# LOCAL REGULATION AND FUNCTION OF IMPORTIN-β1 IN HIPPOCAMPAL NEURONS DURING TRANSCRIPTION-DEPENDENT PLASTICITY

**DOI:** 10.1101/2020.12.02.409078

**Authors:** Yan Jun Lee, Sheeja Navakkode, Chee Fan Tan, Siu Kwan Sze, Sreedharan Sajikumar, Toh Hean Ch’ng

## Abstract

Activity-dependent transcription is critical for the encoding of long-term memories. Regulated nuclear entry of soluble proteins is one method to relay synaptic signals to the nucleus to couple neuronal excitation with transcription. To date, the role of importin-β1 in nuclear shuttling of proteins during activity-dependent transcription has always been inferred but not directly investigated. In this study, we demonstrate activity-dependent nuclear accumulation of importin-β1 from the soma and the synapto-dendritic compartments. Importantly, inhibition of importin-β1 mediated nuclear import during synaptic stimulation impairs long-term plasticity. We show evidence that importin-β1 mRNA-ribosome complex is distributed throughout the synapto-dendritic compartment and synaptic stimulation induces importin-β1 local protein synthesis. Finally, we identified candidate proteins that associate with importin-β1 at the synapse and characterize NDRG1 as an importin-β1 interactor that undergoes activity-dependent translocation into the nucleus. Collectively, our results highlight the crucial role of importin-β1 in the nuclear import of soluble proteins during long-term plasticity.

## INTRODUCTION

Experiments in animal models have established the importance of transcription during different forms of learning (Alberini and Kandel, 2014, Eagle et al., 2016, Yap and Greenberg, 2018). For instance, studies using brain slices and cultured neurons have shown the blocking of transcription during a critical window reduces long-term synaptic potentiation and the efficacy of synaptic transmission (Wiegert et al., 2009, Frey et al., 1996b, Nguyen et al., 1994). Moreover, multiple *in vivo* experiments manipulating transcriptional modulators have resulted in a poorer performance of memory-related tasks in animals (Alberini, 2009, Suzuki et al., 2011, Sekeres et al., 2010). Collectively, these experiments demonstrate the importance of coupling neuronal activation with transcriptional modulation required for inducing persistent changes in the brain during the formation of long-term memories.

To understand how activity-dependent transcription alters neuronal properties, it is not only important to identify how different patterns of activity affect transcription, but also how these electrical or chemical signals are relayed to the nucleus. Mapping out the different mechanisms of signal transduction to the nucleus will shed light on the temporal events that regulate the different waves of activity-driven transcription. Indeed, reports have shown that the several transcriptional waves can be governed by different signalling cascades and the activation of different transcription factors upon neuronal stimulation (Igaz et al., 2002, Wu et al., 2001, Saha et al., 2011, Brigidi et al., 2019). More recently, it has been demonstrated that the neuronal activity patterns, including the duration of activity, can induce multiple waves of transcription, each with its own distinct expression profile (Tyssowski et al., 2018). Critically, the induction of these distinct transcriptional waves requires signalling cascades that trigger the nuclear entry of different soluble proteins (Tyssowski et al., 2018).

There are several mechanisms for proteins to enter the nucleus (Kose et al., 2012, Kimura and Imamoto, 2014, Cautain et al., 2015, Ma et al., 2014a). Apart from the diffusion of small proteins or direct association with the nuclear pore complexes (NPC), many proteins enter the nucleus by binding to nuclear adaptors via its nuclear localization signals (NLS). To date, importins are the most-ubiquitous and best-studied family of the nuclear adaptor proteins (Cautain et al., 2015, Wagstaff and Jans, 2009), and the importin-mediated nuclear import can be categorized into classical and non-classical pathways. In the classical pathway, Imp-α isoforms heterodimerize with Imp-β1 to carry classical NLS-bearing cargo proteins into the nucleus (Stewart, 2007) while the non-classical nuclear import mechanisms engage the Imp-β or β-like importins. There are 22 Imp-β members in humans, of which at least 11 are reported to interact with cargo proteins (Chook and Süel, 2011, Baade and Kehlenbach, 2019, Bourgeois et al., 2020). Imp-β members function independently from Imp-α and most of them enter the nucleus in its monomeric form, with some forming heterodimers with other Imp-β members (Chook and Süel, 2011, Baade and Kehlenbach, 2019). At present, it is unclear if different patterns of activity preferentially recruit specific importins for nuclear entry.

There is increasing evidence showing Imp-β1, through the classical Imp-α/β1 pathway, is engaged in activity driven nuclear entry of plasticity-associated proteins from the soma and the synapto-dendritic compartment (Thompson et al., 2004, Dieterich et al., 2008, Lai et al., 2008, Mikenberg et al., 2007, Dinamarca et al., 2016). For instance, the components of the classical nuclear import pathway, including Imp-β1, is found localized at the synapse and the dendrites, and that they undergo nuclear accumulation in response to activity (Thompson et al., 2004). Moreover, Imp-α is reported to associate with cargo proteins bound for the nucleus in response to activity (Dieterich et al., 2008, Lai et al., 2008, Mikenberg et al., 2007, Dinamarca et al., 2016). However, whether Imp-β1 is truly required for the regulation of gene expression during long-term plasticity has never been directly interrogated. While it appears that Imp-β1’s role in long-term plasticity is indispensable, several questions have been raised to challenge this assumption. First, there is a built-in redundancy in the importin-mediated nuclear import machinery. Many nuclear proteins can associate with different importins and a loss of Imp-β1 function could be compensated by other Imp-β members (Chook and Süel, 2011, Baade and Kehlenbach, 2019). Second, there are non-canonical nuclear import mechanisms that function independently from Imp-β1 (Cautain et al., 2015, Ma et al., 2014b, Oka and Yoneda, 2018, Lever et al., 2015, Kotera et al., 2005). Indeed, for every plasticity-associated protein that engages Imp-β1, there is an equal number of proteins in the activity-driven signalling cascade, such as PKA, Akt, and MEK, that do not have a clear NLS or reported to associate with Imp-β1. Third, the possibility that the nuclear calcium signalling alone is adequate to trigger the first transcriptional wave without the need for importin-mediated nuclear import of soluble proteins has been proposed (Hardingham et al., 2001). Nonetheless, given the spatial origin of the input signal and the kinetics of nuclear import, it is possible that Imp-β1 is crucial for triggering or sustaining distinct transcriptional waves beyond the initial expression of activity-regulated genes.

In this study, we first provide a direct evidence showing the importance of Imp-β1 in activity-driven nuclear signalling and transcription-dependent plasticity. We tested Imp-β1 responsiveness to synaptic stimulation and designed loss-of-function studies using a membrane permeable small molecule inhibitor to block Imp-β1 entry into the nucleus. We report a selective loss of JunB and NFATc3 nuclear entry, a reduction in CREB phosphorylation and the expression of activity-regulated genes, as well as an impairment in hippocampal late-LTP and LTD following blockade of Imp-β1 function during activity. Next, we show that Imp-β1 transcript and protein are found cell-wide, including the synapto-dendritic compartment. Our time-lapse imaging experiments demonstrate that synaptodendritically localized Imp-β1 protein responds to activity by undergoing nuclear translocation while Imp-β1 mRNA undergoes activity-dependent local translation. Finally, we performed co-immunoprecipitation of Imp-β1 coupled with tandem mass spectrometry sequencing and identified a list of candidate proteins that interacted with Imp-β1 in stimulated synaptosomes. We characterized one such protein, NDRG1, that undergoes Imp-β1-mediated activity-dependent nuclear translocation. Taken together, our results confirm the importance of Imp-β1, and more broadly, adaptor protein-mediated nuclear import of soluble proteins, during transcription-dependent plasticity.

## RESULTS

### Neuronal stimulation triggers nuclear entry of Imp-β1 in hippocampal neurons

Previous reports have shown that the classical nuclear import pathway responds to neuronal activity. In particular, Imp-α/β1 complex accumulates in the nucleus upon glutamatergic stimulation (Thompson et al., 2004). To extend these findings, we focused specifically on the role of Imp-β1 in neurons. We tested several antibodies and selected monoclonal antibody (3E9) that showed specificity for Imp-β1 (**Fig. S1A**) and robustly detects the protein in multiple subcellular compartments including the nucleus, dendrites, and the synapses. As expected, 3E9 detects high concentration of Imp-β1 around the nuclear membrane, presumably reflecting the docked Imp-β1 at the NPC (Gorlich et al., 1995, Ström and Weis, 2001).

We next asked if different stimulation paradigms can drive nuclear entry of Imp-β1 in hippocampal neurons. We stimulated cultured hippocampal neurons using a brief pulse of glutamate (40 µM, 5 min) and observed a rapid nuclear accumulation of Imp-β1 in the pyramidal neurons (**Fig. 1A**). Similarly, the depolarization of neurons, in the presence of glycine to preferentially enhance NMDA receptor activation (Corera et al., 2009), also resulted in a robust nuclear accumulation of Imp-β1 (**Fig. 1A**). Importantly, the activity-induced Imp-β1 nuclear translocation is blocked with the addition of APV, an NMDA receptor antagonist (**Fig. 1A**). We also stimulated neurons with bicuculline, a GABA_A_ receptor antagonist, which enhances overall network activity by blocking inhibition and promoting action potential bursting in excitatory neurons (Eisenman et al., 2015). Under these conditions, we observed a modest increase of nuclear Imp-β1 in the pyramidal neurons (**Fig. 1C**). Since importins serve as adaptor proteins that undergo rapid nucleocytoplasmic shuttling, we reasoned that a more synaptic-driven activation via GABAergic inhibition with bicuculline is likely to result in a modest enrichment of Imp-β1 in the nucleus. We also noted that the presence of tetrodotoxin (TTX), which inhibits action potential firing in neurons, did not further reduce Imp-β1 in the nucleus, suggesting a steady state level of nucleocytoplasmic shuttling is maintained in the neurons.

**Figure 1.**
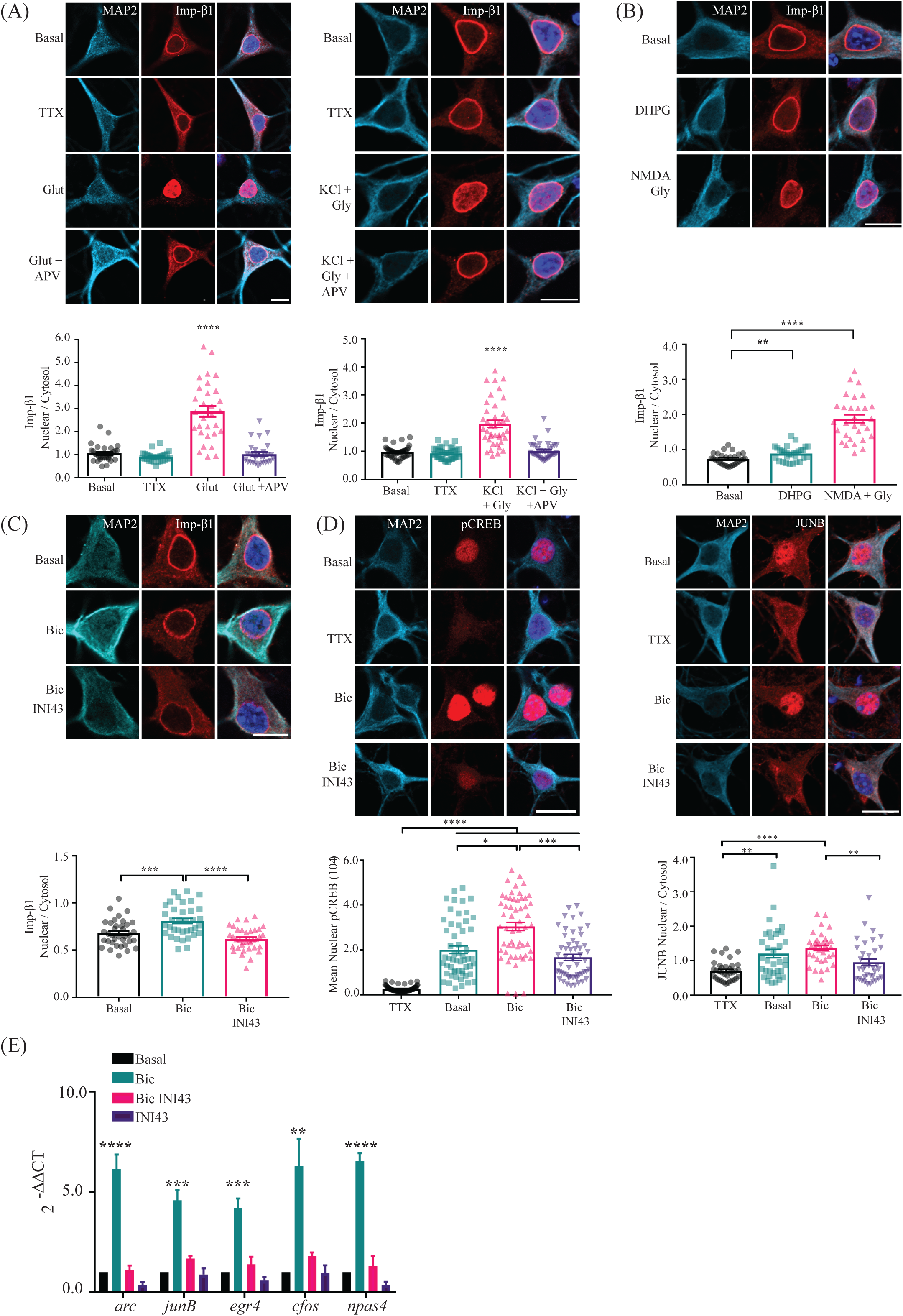
Stimulation-induced nuclear entry of Imp-β1 in hippocampal neurons is critical for nuclear signaling underlying transcription dependent plasticity. **(A)** Cultured mouse hippocampal neurons were treated with either tetrodotoxin (TTX; 1 μM, 1 h), glutamate (Glut; 40 μM, 5 min) or KCl (50 mM, 5 min) + glycine (100 μM, 5 min) in the presence or absence of APV (40 μM, 1 h) before fixing and immunolabeling with antibodies targeted against MAP2 (cyan), Imp-β1 (red) and Hoechst nuclear dye (blue in merged image). The Imp-β1 nuclear-to-cytoplasmic ratio was quantified and group data from independent experiments were plotted. One way-ANOVA with Dunn’s post-hoc analyses were conducted (Glutamate: N=3, KCl: N=2; \*\*\*\**p* < 0.0001). Scale bar, 10 µm. **(B)** Hippocampal neurons were bath applied with DHPG (100 μM, 20 min) or NMDA (20 μM, 5 min) + glycine (20 μM, 5 min) in before fixing and immunolabeling with antibodies targeted against MAP2 (cyan), Imp-β1 (red) and Hoechst nuclear dye (blue in merged image). The Imp-β1 nuclear-to-cytoplasmic ratio was quantified and the group data from independent experiments were plotted. Unpaired T-test analyses were conducted between stimulated and basal conditions (N=2; \*\**p* < 0.01, \*\*\*\**p* < 0.0001). Scale bar, 10 µm. **(C)** Hippocampal neurons were treated with bicuculline (Bic; 40 μM, 30 min) with or without INI43 (5 μM, 1 h) in before fixing and immunolabeling with antibodies targeted against MAP2 (cyan), Imp-β1 (red) and Hoechst nuclear dye (blue in merged image). The Imp-β1 nuclear-to-cytoplasmic ratio was quantified and the group data from independent experiments were plotted. One way-ANOVA with Tukey’s post-hoc analyses (N=2; \*\*\**p* < 0.001, \*\*\*\**p* < 0.0001). Scale bar, 10 µm. **(D)** Representative confocal micrographs of basal and stimulated cultured hippocampal neurons, in the presence or absence of INI43, immunolabeled with antibodies targeted against MAP2 (cyan), pCREB (S133) or JunB (red) and Hoechst nuclear dye (blue in merged image). Cultured neurons were treated with various pharmacological agents including tetrodotoxin (TTX; 1 μM, 1 h), bicuculline (Bic; 50 μM, 30 min) + 4AP (200 μM, 30 min) and INI43 (5 μM, 1 h). For experiments with INI43, neurons were pre-treated with the inhibitor prior to stimulation. The nuclear-to-cytoplasmic ratio of JunB as well as the mean nuclear intensity of pCREB (S133) staining in neurons were quantified and the group data from independent experiments were plotted. Statistical analyses performed on group data use one way-ANOVA with Dunn’s post-hoc analyses (JunB: N=2, pCREB: N=3; \**p* < 0.05, \*\**p* < 0.01, \*\*\**p* < 0.001, \*\*\*\**p* < 0.0001). Scale bar, 10 µm. **(E)** Quantitative PCR (qPCR) analyses on the expression of activity-regulated genes in stimulated cultured hippocampal neurons, with or without INI43 treatment. Cultured neurons were treated with various pharmacological agents including bicuculline (Bic; 50 μM, 30 min) + 4AP (200 μM, 30 min) and INI43 (5 μM, 1 h). For experiments with INI43, neurons were pre-treated with the inhibitor prior to stimulation. Bar graphs depict the 2^-ΔΔCt^ normalized to basal condition. Statistical analyses performed on group data use one way-ANOVA with Tukey’s post-hoc analyses (N=3; \**p* < 0.05, \*\**p* < 0.01, \*\*\**p* < 0.001, \*\*\*\**p* < 0.0001).

To complement the studies in hippocampal cultures, we induced chemical LTP (cLTP) in acute forebrain slices by incubating the slices in Mg^2+^-free artificial cerebrospinal fluid (aCSF) supplemented with forskolin and KCl (Ch’ng et al., 2012). To ensure the slices were sufficiently stimulated, we tracked the increase of the activity-dependent nuclear localization of CRTC1 following cLTP stimulation (Ch’ng et al., 2015). Consistent with the observations made in the hippocampal cultures, Imp-β1 accumulates in the nucleus following cLTP stimulation (**Fig. S1B**). We further investigated if stimulus that trigger other forms of plasticity, specifically long-term depression (LTD), drives Imp-β1 into the nucleus. Both DHPG (100 µM, 20 min) and NMDA (20 µM, 5 min) coupled with glycine (20 µM, 5 min) are known chemical stimuli to initiate LTD in cultures (Waung et al., 2008, Lai et al., 2008). Using both paradigms, we observed a significant nuclear accumulation of Imp-β1 in NMDA- and DHPG-dependent LTD (**Fig. 1B**). Taken together, these experiments confirmed Imp-β1 nuclear entry can be driven by different stimuli, including those associated with long-term potentiation and depression.

### Importin-β1 is critical for transcription-dependent plasticity in hippocampal neurons

To circumvent any deleterious effects of knocking out or knocking down Imp-β1 protein in neurons, we transiently treated the cultured hippocampal neurons with a membrane permeable molecule known as the Inhibitor of Nuclear Import-43 (INI43) during neuronal stimulation. INI43 was identified from an *in silico* screen, using a structure-based approach, for compounds that interfered with Imp-β1-α2-Ran heterodimerization (van der Watt et al., 2016). This molecule has been shown to block the nuclear entry of endogenous Imp-β1 and several of its cargo proteins such as NFAT, NF-κβ and AP-1 complex (van der Watt et al., 2016, Forwood et al., 2001, Stelma and Leaner, 2017). We optimized INI43 treatment parameters in cultured hippocampal neurons and show it blocks Imp-β1 nuclear accumulation during bicuculline and glutamatergic stimulation in pyramidal neurons (**Fig. 1C, S2A and S2B**). We next tested if INI43 can inhibit the nuclear entry of NLS-bearing cargo proteins by selecting NFATc3, a transcription factor that associates with Imp-β1 and responds to activity by undergoing nuclear translocation (Wild et al., 2019, Murphy et al., 2019, Torgerson et al., 1998, Ishiguro et al., 2011). Consistent with published reports, depolarization drives transiently expressed sGFP-NFATc3 into the nucleus and this translocation is partially inhibited by INI43 (**Fig. S2B**). Importantly, we also validated the specificity of INI43 by showing the activity-induced CRTC1 nuclear accumulation in neurons is unaffected by the inhibitor since CRTC1 nuclear entry is independent of Imp-β1 (Ch’ng et al., 2015) (**Fig. S2C**).

Since INI43 inhibits Imp-β1 cargo binding and nuclear entry, we wanted to know the extent of disruption in the activity-driven nuclear signalling pathways following the inhibition. We first looked at the phosphorylation of CREB at Serine 133 (pCREB S133) (Ch’ng et al., 2012, Shaywitz and Greenberg, 1999). Phosphorylation of CREB (S133) depends largely on the activity of different kinases including CamkI, CamkII, PKA and PKC (Johannessen and Moens, 2007). Bicuculline stimulation of cultured hippocampal neurons, in the presence of 4AP to increase the bursting frequency of bicuculline-induced action potential, enhances the phosphorylation of CREB (S133) (Hardingham et al., 2001). However, in the presence of INI43, pCREB S133 is greatly diminished (**Fig. 1D**). Similarly, we asked if the activity-induced nuclear enrichment of JunB, a component of AP-1 complex, is altered with INI43 (Alberini, 2009). As our data indicate, INI43 reduces the JunB nuclear accumulation upon bicuculline/4AP stimulation, suggesting the activity-induced JunB nuclear localization is dependent on Imp-β1 (**Fig. 1D**). Since transcription of many activity-regulated genes rely on the phosphorylation of CREB (Saha et al., 2011), we anticipate that in the absence of stimulus-driven Imp-β1 mediated nuclear import, many of the activity-dependent changes in the nucleus would be impacted. Quantitative PCR (qPCR) analyses of the cultured hippocampal neurons show an increased in the expression of activity-regulated genes namely *arc, junB, egr4*, *cfos* and *npas4* in response to bicuculline/4AP stimulation, and this increment is completely abolished with INI43 (**Fig. 1E**). Critically, the transcript level of the activity-independent housekeeping gene, HPRT1, in the differentially treated samples are comparable across all conditions, indicating the cells are viable throughout the treatment (**Fig. S1D-i**). Moreover, the immunoreactivity of cleaved caspase 3, a hallmark for apoptosis, is not increased in INI43 treated neurons and are significantly lower than H_2_O_2_ induced apoptotic neurons (**Fig. S1D-ii;** (Park et al., 2016). This further indicates that the transient exposure of INI43 does not trigger apoptosis in neurons and the cells are viable throughout the experiments. Collectively, our data indicate Imp-β1 mediated nuclear import mechanism is required for initiating nuclear events underlying long-term plasticity.

Late phase LTP (late-LTP) in the CA3 to CA1 hippocampal circuit is transcription-dependent (Nguyen and Kandel, 1997). Experiments have shown the addition of actinomycin D and other transcriptional inhibitors impairs late-LTP (Frey et al., 1996a, Nguyen et al., 1994). To evaluate if Imp-β1 mediated nuclear entry is also necessary for transcription-dependent late-LTP and/or late-LTD, we employed a two-pathway model (**Fig. 2A)** using *in vitro* electrophysiology. Bath application of INI43 for 1 h after a stable baseline of 30 min in wild-type mouse hippocampal slices did not alter the baseline recordings (**Fig. 2B**). The slope of the fEPSPs evoked every 5 min after drug application remained unchanged for a total of 180 min. No significant changes were observed in fEPSP values throughout the recording when compared with its own baseline (*p* > 0.05, Wilcox test). The results confirm that the basal synaptic transmission is not affected when Imp-β1 mediated nuclear transport is blocked by INI43.

**Figure 2.**
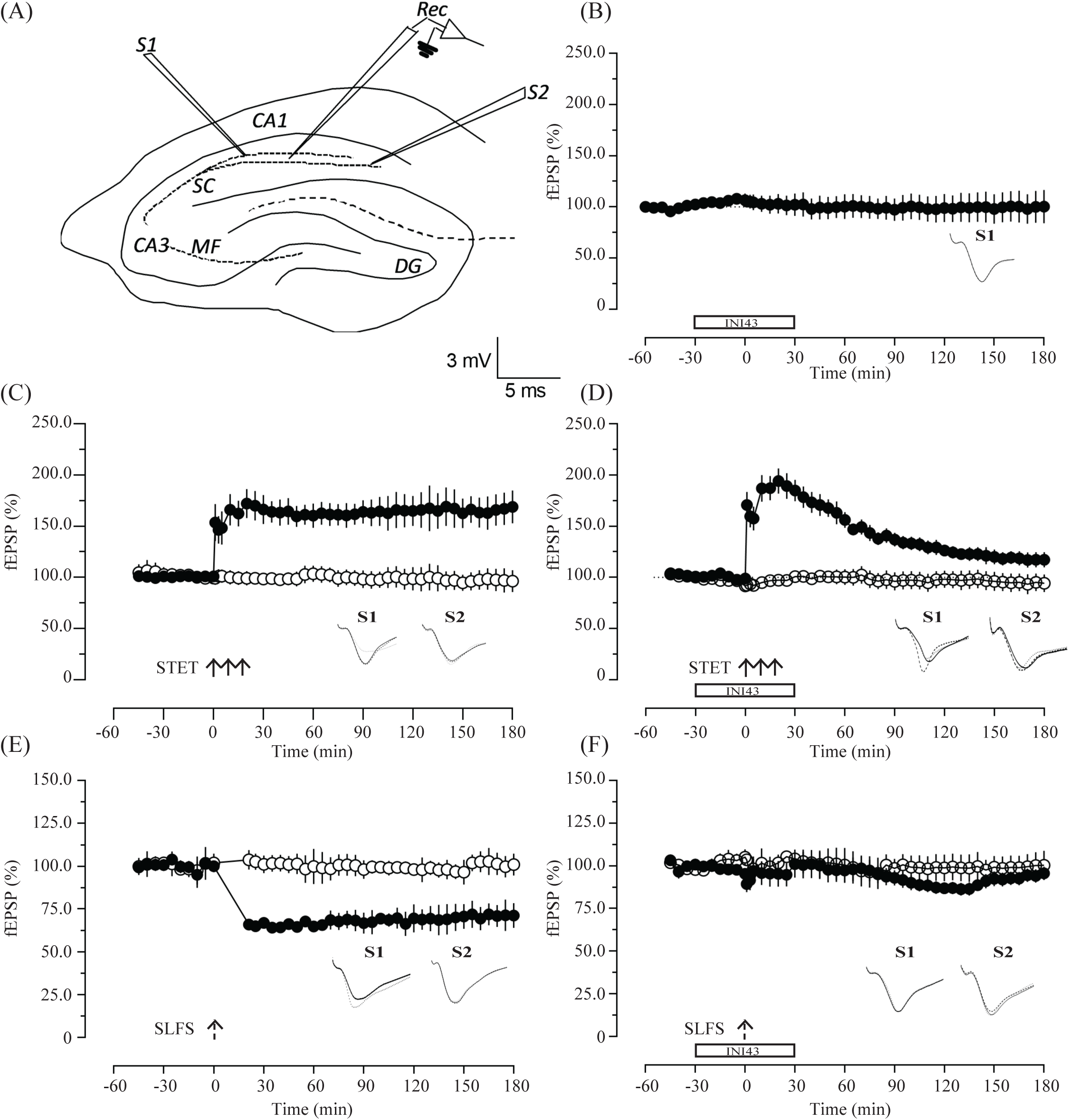
Inhibition of Imp-β1 prevents late-LTP and LTD. **(A)** Schematic representation of a transverse hippocampal slices showing the positioning of electrodes in the CA1 region for field EPSP (fEPSP) recording. Two stimulating electrodes, S1 and S2, were placed in the stratum radiatum to stimulate two independent Schaffer Collateral synaptic inputs to the same neuronal population. Recording electrode (Rec) was placed in the CA1 apical dendrites between the two stimulating electrodes. Calibration scale bars for all analog traces: vertical: 3 mV horizontal: 5 ms. **(B)** Basal synaptic transmission is not affected by the application of INI43 (5 μM). Application of INI43 for 1 h after recording a stable baseline for 30 min did not alter the baseline values and it remained stable for 3 h. (n=6) **(C)** Late-LTP was induced by STET (arrows) in S1 (filled circles) after recording a stable baseline for 30 min. Open circles represent control input S2, which remained stable throughout the 3 h period of recording. The test stimulation was given every 5 min. (n=7) **(D)** Experimental design was the same as in **(C)**, but with the exception that INI43 (5 μM) was applied 30 min before and 30 min after the induction of late-LTP, which resulted only in an early-LTP. (n=7) **(E)** L-LTD was induced using a SLFS after recording a stable baseline for 30 min which resulted in a depression that was stable for 3 h recording time. (n=8) **(F)** Experimental design same as in **(E)** but INI43 (5 μM) was applied 30 min before and 30 min after the induction of L-LTD in S1, that resulted in the total abolishment of LTD (n=8). Control inputs S2 were stable in both experiments throughout the period of recording and the data were collected with at least three independent biological replicates.

As a control experiment, we showed that the application of a strong tetanus (STET) that consisted of 100 Hz, 100 pulses at 0.2 ms resulted in a late-LTP that was stable for a recorded period of 180 min (**Fig. 2C**). A statistically significant potentiation was observed immediately after tetanus at 1 min (*p* = 0.0156, Wilcox test; *p* = 0.0006, U-test) and it persisted until 180 min (*p* = 0.0156, Wilcox test; *p* = 0.0006, U-test). The synaptic potentiation was input specific as no significant shift in fEPSPs in the unpaired control input S2 was observed (open circles). Next, to test the role of Imp-β1 in transcription-dependent plasticity, late-LTP was induced using STET in the presence of INI43 (**Fig. 2D**). After recording a stable baseline for 30 min, INI43 was bath applied to the hippocampal slices for 1 h and late-LTP was induced 30 min after the beginning of the drug application in S1. Induction of late-LTP during the inhibition of Imp-β1 mediated nuclear import by INI43 prevented the late-LTP without affecting the persistence of early-LTP. A significant potentiation was observed immediately after tetanus at 1 min (*p* = 0.0156, Wilcox test; *p* = 0.0006, U-test), and it remained significant until 120 min (*p* = 0.0156, Wilcox test; *p* = 0.0262, U-test), which then decayed back to baseline at 180 min (*p* = 0.973, Wilcox test) (**Fig. 2D;** filled circles). No significant changes were observed in the control input S2 and it remained stable throughout the period of recording (open circles). When comparing between the two groups of LTP with and without INI43 (**Fig. 2C & 2D)**, no significant difference was observed from 1 min until 60 min (at 1 min, *p* = 0.2086, U-test; at 60 min, *p* = 0.7104, U-test) but from 120 min onwards, it showed significant difference and remained significant until 180 min (at 120 min, *p* = 0.0012, U-test; at 180 min *p* = 0.0023, U-test).

Since importins are released during NMDA receptor activation and that LTD also involves NMDA-receptor activation, we tested if Imp-β1 mediated transcriptional mechanisms play a role during LTD. Late-LTD was induced using a strong low frequency stimulation (SLFS) consisted of 900 pulses (Sajikumar and Frey, 2003, Shetty et al., 2015), and a robust synaptic depression was recorded for up to 180 min post-SLFS (**Fig. 2E**). Application of SLFS led to a significant decrease in fEPSP values from 1 min (*p* = 0.0313, Wilcox test; *p* = 0.0022, U-test) and it remained stable until the end of the recording at 180 min (*p* = 0.0313, Wilcox test; *p* = 0.0022, U-test). Subsequently, to study the role Imp-β1, we applied INI43 during the induction of late-LTD. A stable baseline was recorded for 30 min and a late-LTD was induced in S1 using SLFS. We repeated the same experiment as in **Fig.2E** but in the presence of INI43, 30 min before and after SLFS (**Fig. 2F**). Interestingly, unlike LTP, we found that the LTD was significantly impaired at an earlier time point. Field EPSP values were significantly different at 1 min (*p* = 0.0391, Wilcox test; *p* = 0.0112, U-test) and then from 5 min onwards, it was not significant (*p* = 0.7422, Wilcox test; *p* > 0.9999, U-test) until the end of the recorded period of 180 min (*p* > 0.9999, Wilcox test; *p* = 0.7416, U-test). The control input S2 did not show significant changes in both sets of experiments for the entire recording period (open circles). When comparing between the two groups of LTD with and without INI43 (**Fig. 2E & 2F**), we observed a significant difference from 1 min (*p* = 0.008, U-test) until 180 min (*p* = 0.0027, U-test).

Altogether, the hippocampal slice recordings further reinstated the importance of Imp-β1 in transcription-dependent plasticity by directly showing the inhibition of Imp-β1 mediated nuclear import negatively affected the long-term plasticity. Specifically, we showed the blockade of Imp-β1’s function impaired late-LTP while leaving the early-LTP unaffected in tetanized hippocampal slices. In addition, our data suggest that the Imp-β1 mediated nuclear import could be critical for late-LTD and the reasons for the impairment during early LTD is revisited in discussion.

### Importin-β1 is localized at the synapto-dendritic compartment and undergoes activity-dependent nuclear translocation

Having established the importance of Imp-β1-mediated nuclear import during transcription-dependent plasticity, we next explored the role and regulation of Imp-β1 in the synapto-dendritic compartment. Imp-β1 is ubiquitously detected cell-wide and to verify endogenous localization of Imp-β1 in the synapto-dendritic compartment, we virally expressed GFP to label spines in excitatory neurons (Thompson et al., 2004). Imp-β1 fluorescent puncta are found throughout the synapto-dendritic compartment including a large majority of GFP-positive spines (**Fig. 3A**). To examine if the synapto-dendritic localized Imp-β1 undergoes activity-dependent movement into the nucleus, we transiently expressed Imp-β1-Dendra2 or Dendra2 alone constructs in excitatory neurons using a CamKIIα promoter for a short period (14-16 h) to limit overexpression and performed time-lapse imaging. To label a subpopulation of Imp-β1, we photoconverted the Dendra2 signal in three distinct synapto-dendritic compartments (R1-R3) and measured the nuclear accumulation of the photoconverted Dendra2 signal in the nucleus in the presence or absence of bicuculline stimulation. Importantly, each photoactivated region showed an increase in Dendra2 (red-shifted) signal immediately upon photoconversion without a corresponding increase at the soma (**Fig. 3B**). Time-lapse images obtained over 15 min (1 min / frame) indicate a gradual (starting at 5 min post-conversion) but significant nuclear accumulation of the photoconverted Imp-β1-dendra2 signal in the bicuculline stimulated neurons (**Fig. 3C**). Unlike other synapse-to-nucleus proteins that show increasing nuclear accumulation, the increase in nuclear Imp-β1 reaches a plateau and achieves equilibrium which is consistent with the idea that Imp-β1 undergoes rapid nucleocytoplasmic shuttling. Notably, we did not observe a significant nuclear accumulation of the photoconverted Imp-β1-dendra2 in basal neurons as well as in neurons expressing the Dendra2 alone construct (**Fig. 3C & S3**). The appearance of the photoconverted Dendra2 control signal in the nucleus shortly after photoconversion is likely to be due to diffusion and is independent of bicuculline stimulation. Taken together, our live cell experiments illustrate the activity-dependent nuclear translocation of synapto-dendritic Imp-β1.

**Figure 3.**
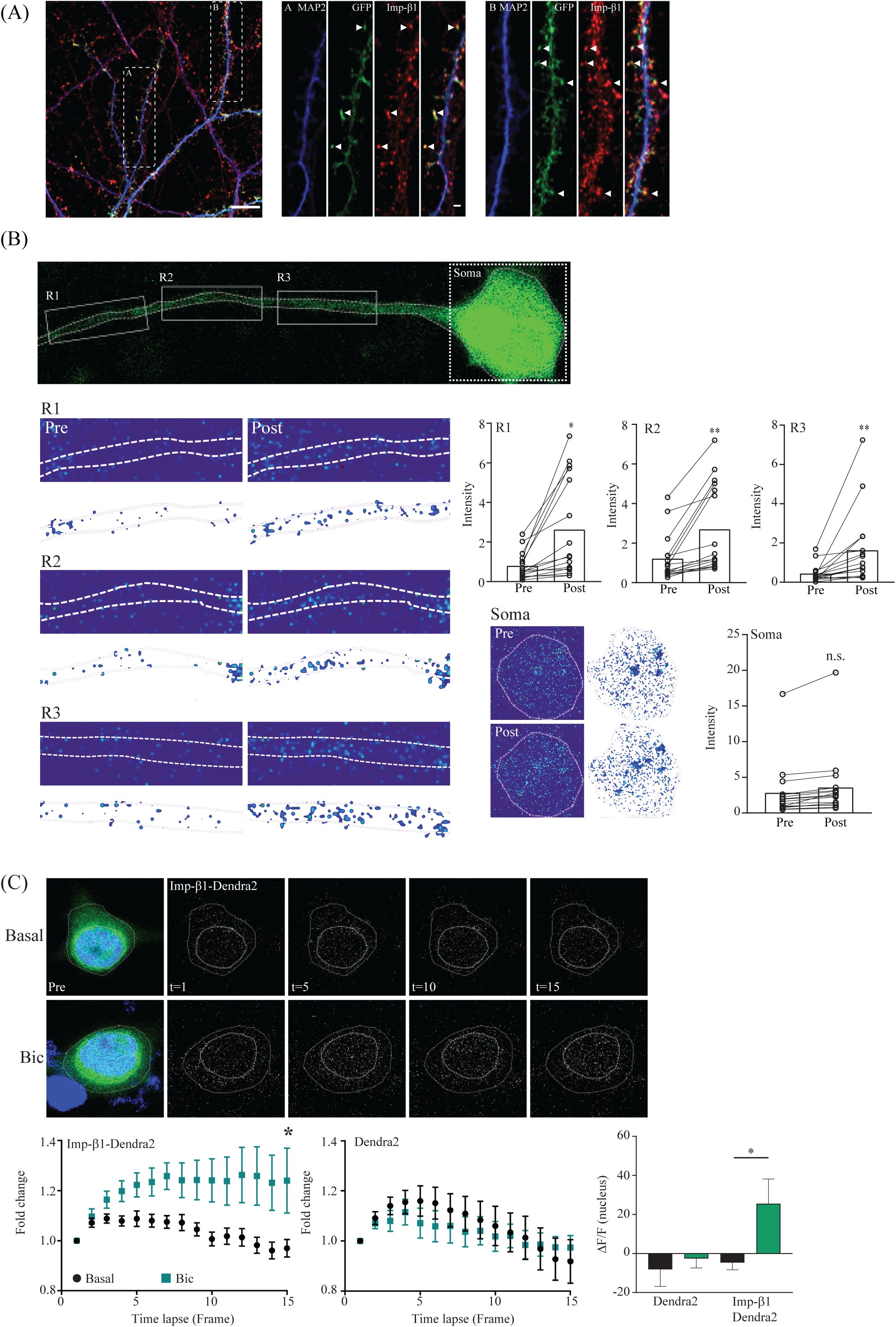
Imp-β1 is localized at the synapto-dendritic compartment and undergoes activity dependent nuclear translocation. **(A)** Cultured mouse hippocampal neurons expressing virally delivered GFP driven by CamKIIα promoter. Neurons were fixed and immunolabeled with antibodies targeted against MAP2 (blue), GFP (green) and Imp-β1 (red). White dotted boxes (A & B) are magnified with arrows indicating Imp-β1 positive spines. Scale bar 10 µm (1 µm in magnified images). **(B)** Cultured rat hippocampal neurons expressing Imp-β1-Dendra2 or Dendra2 alone constructs were photoconverted at three regions indicated by the white boxes (R1-R3) using UV (405 nm) laser and the nuclear accumulation of the photoconverted signal (568 nm) was tracked for 15 min (at 1 frame per min). Representative confocal images indicating the pre- and post-photoconverted regions in dendrites and soma with the corresponding before-after graphs of the red-shifted Dendra2 (568 nm) intensity in the photoconverted region. Statistical analyses performed on group data use Mann-Whitney non-parametric t-test (N=3, n=16; \**p* < 0.05, \*\**p* < 0.01). **(C)** Time lapse images of the somatic photoconverted Imp-β1-Dendra2 signals (every 5^th^ frame) are shown for basal and bicuculline (40 µM) stimulated neurons. The line graphs show normalized photoconverted signal in the nucleus. Statistical analyses performed on group data use Mann-Whitney non-parametric T-test for comparing the same time point between treatment groups and one way-ANOVA with Dunn’s post-hoc analyses to compare the different time points against t=1 within the same treatment group. (N=3, Basal: Dendra2, n=7; Bic: Dendra2, n=5; Basal: Imp-β1-Dendra2, n=10; Bic: Imp-β1-Dendra2, n=11, \**p* < 0.05, Bic-Imp-β1-Dendra2, t=1 as compared to t=3 to 13; \**p* < 0.05; Bic-Imp-β1-Dendra2 as compared to Basal-Imp-β1-Dendra2, from t=5 to 15 \**p* < 0.05). Bar graph depicts the change in red shifted Dendra2 (568 nm) nuclear intensity between the first and the final time points (t=1 and t=15; \**p* < 0.05).

### Importin-β1 is locally translated in the synapto-dendritic compartment

Since Imp-β1 translocates from the synapto-dendritic compartment to the nucleus, it is likely that the cargo binding of Imp-β1 is locally regulated at or near the synapse. Unlike Imp-α, where it has been shown to tether to the cytoplasmic tail of the NR1-1a subunit of the NMDA receptors and is released during glutamatergic stimulation (Jeffrey et al., 2009), nothing is known about the regulation of Imp-β1 at synapses. Studies have reported the presence of Imp-β1 transcripts in mid-axons, dendrites and in the presynaptic terminals (Hafner et al., 2019, Perry et al., 2012b, Poon et al., 2006), suggesting that Imp-β1 may be regulated through activity-dependent local translation in the synapto-dendritic compartment (Tushev et al., 2018, Biever et al., 2019, Holt et al., 2019).

Previously, two Imp-β1 isoforms have been reported in neurons, with the longer isoform preferentially targeted to axons (Perry et al., 2012a). Using algorithms to detect the differential usage of PolyA signals (Proudfoot, 2011), we identified additional PolyA signals in the mouse Imp-β1 mRNA (NM_008379.3) that would correspond to a full-length Imp-β1 mRNA (5.9 kb), as well as the long and short isoforms as previously described by Perry *et al*. (2012a). To detect the relative abundance of these different Imp-β1 mRNA isoforms, we designed primers targeting different regions of the Imp-β1-3’UTR, specifically to identify the presence of the full-length (F1 and F2), the long (L1 and L2) and the short (S1) mRNA isoforms and performed qPCR on purified synaptosomes extracted from mouse forebrain (**Fig. 4A**). The qPCR assay detected all three Imp-β1 mRNA isoforms, with the full-length and combined (full-length and long) isoforms representing 5.1 % and 18.6 % respectively (**Fig. 4A**). To further distinguish if a specific Imp-β1 isoform is enriched in the synaptosomes, we extracted and profiled the mRNA from the forebrain cytosolic fraction and observed a similar distribution for all three Imp-β1 isoforms as seen in the synaptosome preparation, suggesting no specific enrichment of any one type of mRNA isoform at the synapse (**Fig. 4A**). Importantly, the same results were observed using either random hexamers or oligo-dT priming to generate cDNA from the total synaptosome RNAs pool and all primer pairs selected for this study are specific for each isoform with no detectable contamination (**Fig. 4A**).

**Figure 4.**
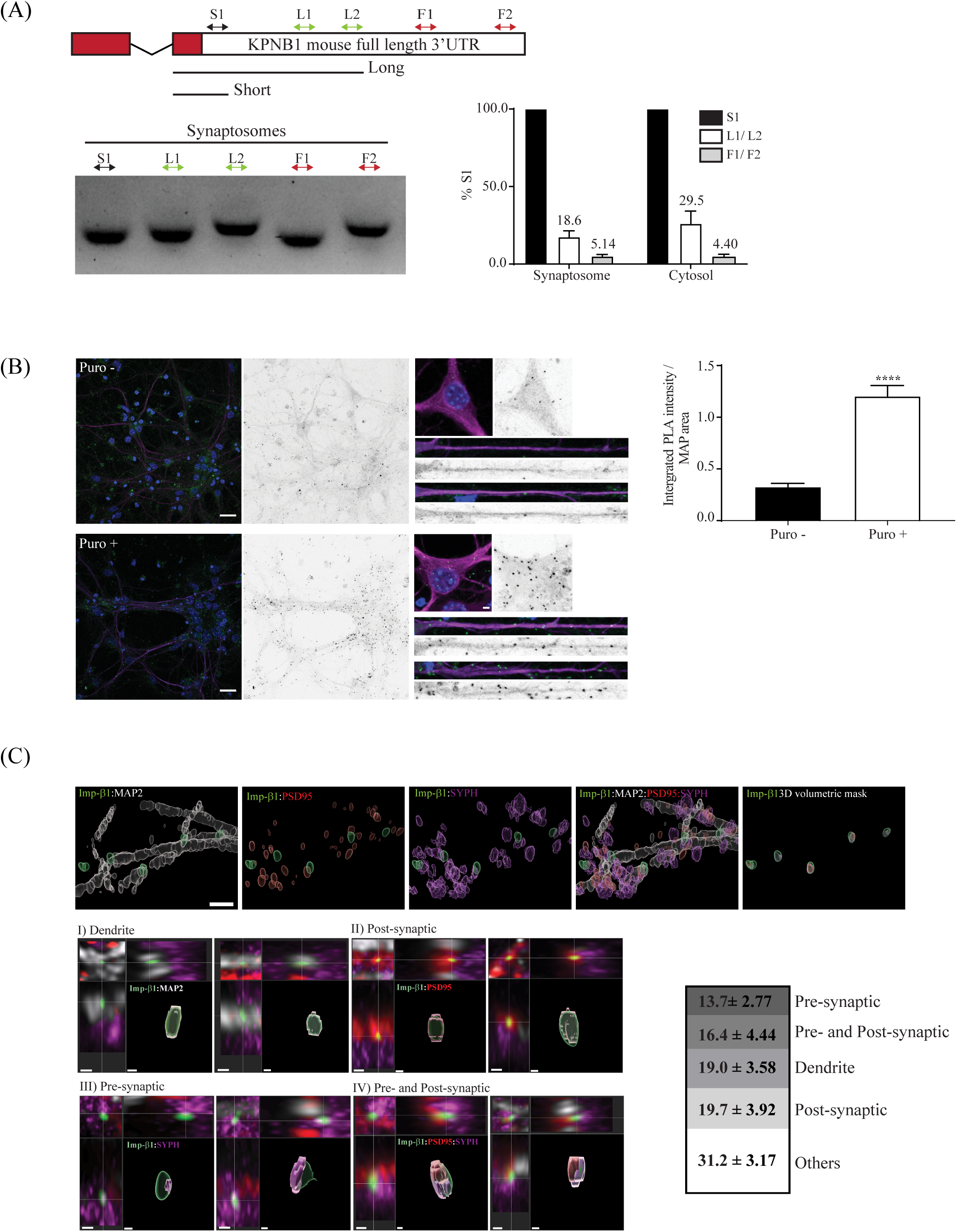
Imp-β1 is locally translated in the synapto-dendritic compartment basally. **(A)** Schematic of the mouse Imp-β1 mRNA showing the last two exons followed by the full length 3’UTR. The red boxes indicate exons in the open reading frame (ORF) while the white box indicates the full length 3’ UTR with the relative size of the long and short 3’UTR isoforms indicated as lines. The bidirectional arrows indicate the relative location of primers used to detect full-length isoform (F1 and F2), long isoform (L1 and L2), and short isoform (S1). The qPCR products were separated on agarose gel and the qPCR results were represented in bar graph depicts as the fold change normalized to the common primer (S1), obtained from both the cytosolic fraction and the purified synaptosomes. The percentage of each splice isoform relative to S1 is indicated at the top of the bar graph (N=3). Note that L1/L2 values include long and full-length isoforms. **(B)** Cultured mouse hippocampal neurons were pre-treated with cycloheximide (CHX; 355 µM, 30min) before Puro-PLA, in the presence or absence of puromycin (Puro; 3 µM, 15 min). Cells were fixed and immunolabeled with antibodies targeted against MAP2 (violet) and Hoechst nuclear dye (blue in merged image). The puro-PLA signal specific for Imp-β1 appears as green puncta (inverted to black-white for contrast) and is localized in soma and synapto-dendritic compartment. The integrated Puro-PLA intensity normalized to total MAP2 area in the field was quantified and the group data from independent experiments were plotted. Unpaired T-test analyses were conducted between the conditions with and without puromycin (N=3; n=30; \*\*\*\**p* < 0.0001). Scale bar, 2 µm. **(C)** Cultured mouse hippocampal neurons were pre-treated with cycloheximide (CHX; 355 µM, 30 min), before undergoing Puro-PLA (green) in the presence of puromycin (Puro; 3 µM, 15 min), followed by fixation and immunolabeling with antibodies targeted against MAP2 (grey), PSD95 (red) and synaptophysin (Syph; violet). Neurons were imaged and deconvoluted using Zeiss AiryScan deconvolution module and signals for each channel were rendered into a three-dimensional object and analyzed for colocalization with Imp-β1 Puro-PLA signal using IMARIS (*Upper panels*). Individual Imp-β1-puro-PLA puncta that colocalized with synapto-dendritic markers are projected along their 3 axes (*xy, xz* and *yz*) and categorized as dendrites (MAP2; Category I), post-synaptic (PSD95; Category II), pre-synaptic (SYPH; Category III), and pre- and post-synaptic (PSD95 and SYPH; Category IV) compartments (*lower panels*). The relative distribution for each category is indicated in the bar chart (N=3, n=161). Scale bar, 10 µm (*Upper panels*) or 2 µm (*lower panels*).

To demonstrate if Imp-β1 is locally translated, we performed puromycin labeled proximity-ligation assay (Puro-PLA) in cultured hippocampal neurons. Puro-PLA enables the visualization of newly synthesized Imp-β1 proteins by detecting puromycylated nascent protein chains at the site of synthesis. The subsequent spatial coincidence between the antibodies targeted against puromycin and Imp-β1 will produce a fluorescent signal that identifies newly translated Imp-β1 proteins. We pre-incubated the cultured neurons with cycloheximide (CHX) prior to puromycin labeling to stall the ribosome and immobilize the nascent chain to the ribosomal machinery, thereby allowing the detection of the entire ribopuromycylated structure at the site of synthesis (**Fig. S4A-i**) (David et al., 2012, tom Dieck et al., 2015). To examine the specificity of the labeling and to establish baseline signals, we optimized the assay by incubating with single probe (plus or minus only; **Fig. S4A-ii**) or in the absence of puromycin (**Fig. 4B**) to demonstrate low background. In contrast to the controls, the complete protocol with both plus and minus probes detected robust signal for Imp-β1-ribosome complex not only in the neuronal soma, but also in the MAP2 positive dendrites (**Fig. 4B and S4A-ii**).

Since Imp-β1 is locally translated along the proximal and distal segments of the dendrites, we wanted to further classify these ribopuromycylated structures in the synapses. To get a quantitative measure, we triple labeled all samples with antibodies targeted against the pre-synaptic marker synaptophysin (SYPH), post-synaptic marker PSD95, along with MAP2 to identify dendrites. To resolve the three-dimensional structures of the synapse along with the colocalization of Imp-β1 Puro-PLA signal, we performed confocal imaging using AiryScan detector followed by image deconvolution and volumetric reconstruction of the synapse with IMARIS. We rendered three-dimensional surfaces based on the Imp-β1 Puro-PLA signal and examined the extent of colocalization between Imp-β1 Puro-PLA signal and all the synaptic and dendritic markers (**Fig. 4C**). Most of the Imp-β1 Puro-PLA structures were classified into one of the four specific categories: First, approximately 19.0 % of the analysed Imp-β1 Puro-PLA structure shared at least partial colocalization with MAP2 dendritic marker alone, indicating the presence of ribosome-bound Imp-β1 translational complex in the dendrites (**Fig. 4C**). We next analysed the 3D-structures that colocalized with the synaptic markers and observed a proportion of them exhibiting at least a partial overlap with either the post-synaptic marker (19.7 %), the pre-synaptic marker (13.7 %) or with both synaptic markers (16.4 %). Importantly, a subset of these volumetric structures is localized close to or overlapping with MAP2 signal, indicating the proximity to the dendrites (**Fig. 4C**). Finally, almost one-third (31.7 %) of the Imp-β1 Puro-PLA structures are found in surrounding glial cells (**Fig. 4C**). This is not unexpected since Imp-β1 protein expression is found in both the glia and the neurons. Collectively, our assays reveal the presence of Imp-β1 mRNA in the synaptosomes and local translation of Imp-β1 in the synapto-dendritic compartment.

### Synaptic stimulation enhances local translation of importin-β1 in synapto-dendritic compartment

We then proceeded to determine if neuronal activity and synaptic stimuli can enhance Imp-β1 local translation in the synapto-dendritic compartment. Stimulation of biochemically purified synaptosomes has been reported to increase CaMKIIα and BDNF translation (Bagni et al., 2000, Gharami and Das, 2014). Using a similar approach, we purified synaptosomes from adult mouse forebrain and stimulated it with KCl in the presence of glycine, and performed affinity purification to enrich for Imp-β1 protein (Corera et al., 2009). As predicted, we consistently observed an average of 1.5-fold increment in Imp-β1 expression in stimulated synaptosomes as compared to those with mock treatment (**Fig. 5A-i**). Notably, the newly synthesized Imp-β1 is lost when synaptosomes were stimulated in the presence of a protein synthesis inhibitor, cycloheximide (CHX; **Fig. 5A-ii**).

**Figure 5.**
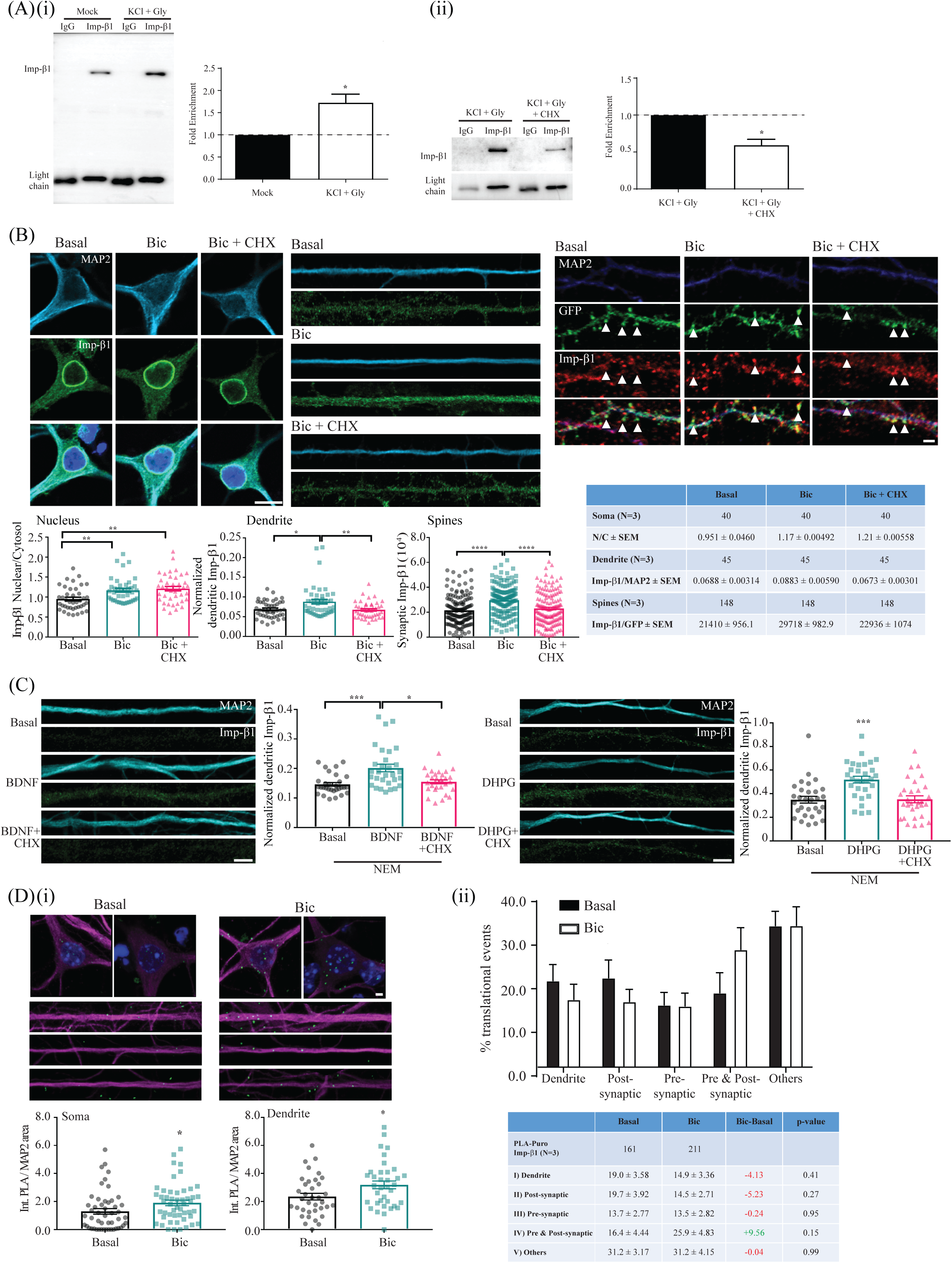
Synaptic stimulation enhances local translation of Imp-β1 in synapto-dendritic compartment. **(A-i)** Synaptosomes purified from adult male mice forebrain were either left untreated (mock) or stimulated with KCl (50 mM, 5 min) and glycine (100 μM, 5 min), followed by co-IP using Imp-β1 antibody (3E9). Blots were probed with 3E9 and band intensities were quantified (N=5). Unpaired T-test analysis was conducted between stimulated and basal condition. \**p* < 0.05. **(A-ii)** Experiment performed as described in (**A-i**) for stimulation, with or without cycloheximide (CHX; 50 μM, 30 min prior and during KCl treatment; N=3). Unpaired T-test analysis was conducted between stimulated synaptosomes with and without CHX treatment. \**p* < 0.05. **(B)** Representative confocal micrographs of basal and bicuculline (Bic; 40 μM, 30 min) stimulated cultured hippocampal neurons, with or without cycloheximide (CHX; 50 μM, 30 min) treatment. The neurons were then immunolabeled with antibodies targeted against MAP2 (cyan), Imp-β1 (green) and Hoechst nuclear dye (blue in merged image). Imp-β1 intensities were quantified for the nucleus (nuclear / cytoplasmic) and dendrites (MAP2 normalized). Group data is represented as bar plots and in the table. Statistical analyses performed on group data use one way-ANOVA with Dunn’s post-hoc analyses (\**p* <0.05, \*\**p* <0.01). Scale bar, 10 µm. For spines, neurons expressing GFP as described in Fig. 3A. were fixed and immunolabeled with antibodies targeted against MAP2 (blue), GFP (green) and Imp-β1 (red). The synaptic Imp-β1 staining were quantified and the group data is represented as bar plots and indicated in the table. Statistical analyses performed on group data use one way-ANOVA with Dunn’s post-hoc analyses (\*\*\*\**p* < 0.0001). Scale bar, 2 µm. **(C)** Representative confocal micrographs of basal, BDNF- (50 ng/mL, 20 min) and DHPG- (100 μM, 5 min with 15 min recovery) treated cultured hippocampal neurons, with or without cycloheximide (CHX; 50 μM, 30 min) prior to and during stimulation. All neurons were also treated with NEM (50 µM, 10 min) and MG132 (5 µM, 30 min) before and during stimulation. The neurons were then immunolabeled with antibodies targeted against MAP2 (cyan) and Imp-β1 (green). The normalized dendritic Imp-β1 staining to MAP2 staining were quantified and the group data is represented as bar plots. Statistical analyses performed on group data use one way-ANOVA with Dunn’s post-hoc analyses (N=2, \**p* <0.05, \*\*\**p* < 0.001). Scale bar, 10 µm. **(D-i)** Cultured mouse hippocampal neurons in basal and bicuculline (Bic, 40μM, 30 min) stimulated states were pre-treated with cycloheximide (CHX; 355 µM, 30 min) followed by Puro-PLA in the presence of puromycin (Puro; 3 µM, 15 min) before being fixed and immunolabeled with antibodies targeted against MAP2 (violet) and Hoechst nuclear dye (blue in merged image). The Imp-β1 Puro-PLA signal is green. The integrated Puro-PLA intensity normalised to total MAP2 area of soma and dendrites were quantified and the group data from independent experiments were plotted. Unpaired T-test analyses were conducted between basal and stimulated groups (N=3, \**p* < 0.05). Scale bar, 2 µm. **(D-ii)** Cultured mouse hippocampal neurons in basal and bicuculline (Bic, 40μM, 30 min) stimulated states were treated, processed, and analysed as described in Fig. 4C. The percentage of each category in basal and stimulated states were quantified and shown in bar graph and in the table.

As synaptosomes contain both the pre- and the post-synaptic structures, we were unable to determine whether one or both the subcellular compartments undergo local translation of Imp-β1. We wanted to identify if the increase in Imp-β1 local translation could be detected in the synapto-dendritic compartment of the stimulated neurons. Unlike bath application of KCl or glutamate which resulted in a robust Imp-β1 nuclear accumulation (**Fig. 1A**) accompanied with a significant reduction in dendritic Imp-β1 (**Fig. S5A**), we observed a moderate Imp-β1 nuclear accumulation following bicuculline-induced action potential bursting (**Fig. 1C**). We reasoned that under these circumstances, the effects of local translation could be observed during the initial stages of bicuculline stimulation. We briefly stimulated the hippocampal neurons with bicuculline (40 µM, 30 min) and showed nuclear accumulation of Imp-β1 that is not sensitive to CHX treatment (**Fig. 5B**). We then quantified dendritic localized Imp-β1 as well as in the GFP-labeled spines and observed a significant increase in Imp-β1 staining in both the subcellular compartments (**Fig. 5B**). Of note, unlike in the nucleus, the increase in the dendritic and post-synaptic Imp-β1 were abolished after CHX treatment (**Fig. 5B**).

To rule out the possibility that the activity-driven increment of synapto-dendritic Imp-β1 is due to the transport of newly-synthesized protein from the soma, we employed N-ethylmaleimide (NEM), a small molecule inhibitor commonly used to inhibit ATP-driven molecular motors including kinesin, dynein and myosin (Pfister et al., 1989). The addition of NEM in cultured hippocampal neurons does not only block bidirectional vesicular trafficking in axons, it also completely abolishes the motor-dependent movement of cytosolic cargo proteins (Scott et al., 2011). To further optimize NEM concentration in neurons, we tracked lysosome movement using lysotracker as a proxy for active transport since lysosome trafficking engages both the dynein and kinesin motor proteins for its bidirectional transport in neurons (Goo et al., 2017, Pu et al., 2016). We found that a brief incubation of NEM (50 µM; 10 min) in cultured neurons effectively blocks the bidirectional displacement of lysosomes, with a significant reduction in the average displacement of lysosomes in the NEM-treated culture compared to mock treatment (**Fig. S5B**). Individual traces of the most actively moving lysosomes within each field show NEM treatment severely impacted its cumulative displacement (Mock: 13.3 µm ± 5.4; NEM: 3.3 µm ± 1.6; **Fig. S5B**).

At this NEM concentration, most of the active transport in the neuronal processes is halted without substantially impacting the translational machinery as NEM inhibits translation only in the millimolar range (Hatey and Gaye, 1978, Wu and Ibrahim, 1980). To stimulate local translation in NEM pre-treated neurons, we used BDNF, a well-studied neurotrophic factor that regulates different forms of neuronal plasticity (Kuipers et al., 2016, Santos et al., 2010). For instance, the application of BDNF in hippocampal neurons enhances glutamatergic synaptic transmission, which in turns, increases spontaneous firing rate and excitatory post-synaptic current. More importantly, BDNF-induced plasticity requires local protein synthesis in dendrites (Kang and Schuman, 1996). The bath application of BDNF (50 ng/mL; 20 min) on NEM pre-treated neurons resulted in a significant increase in dendritic Imp-β1 signal, which is abolished in the presence of CHX (**Fig. 5C**). To complement BDNF stimulation, we used another small molecule agonist, DHPG, that is known to bind and activate mGluR1/5 receptors to trigger mGluRs-LTD (Huber et al., 2000, Fitzjohn et al., 1999). Like BNDF, DHPG-mediated mGluRs-LTD is rapid and requires local protein synthesis (Waung et al., 2008). Similar to BDNF results, bath application of DHPG (100 µM, 5 min) in NEM pre-treated neurons resulted in an increase in dendritic Imp-β1 signal, which is eliminated in the presence of CHX (**Fig. 5C**). Importantly, neither NEM treatment nor BDNF or DHPG bath application altered dendritic MAP2 protein level that is used for normalization (**Fig. S5C**).

If neuronal activity drives local translation of synapto-dendritic Imp-β1, we predict that there will be an increase in the recruitment of its ribosomal translation machinery in the dendrites and/or synapses. To that end, we stimulated neurons with bicuculline (40 µM, 30 min) and immobilized the ribosomes with their nascent peptides in the presence of CHX before performing Puro-PLA. We observed an activity-dependent increase in the integrated signal of Imp-β1 Puro-PLA signal both in the soma and in the dendrites (**Fig. 5D-i**). Next, we asked if the increase in the neuronal activity alters the subcellular distribution of these Imp-β1 Puro-PLA structures. We anticipate that the activity-dependent increase of Imp-β1-ribosomal complex in the synapto-dendritic compartment to be disproportionately localized near or at synaptic sites (Rangaraju et al., 2017). Using the same three-dimensional classification scheme previously described in **Fig. 4C**, we mapped all the Imp-β1 Puro-PLA structures and its colocalization with the pre-post synaptic markers in the bicuculline-stimulated neurons and compared that against neurons under basal condition. Results indicate a trend towards an increase in Imp-β1 Puro-PLA structures that colocalized with both the pre- and post-synaptic markers (Category *IV*) in the bicuculline-treated samples versus those in the basal condition (**Fig. 5D-ii**). In contrast, the distribution of Imp-β1 Puro-PLA structures is reduced in all other categories. These suggest the possibility that activity drives an increase in Imp-β1 local translation close to or at the synapses. While we observed a moderate increase in Imp-β1-ribosome structures, the bicuculline-induced action potential bursting affects only a subset of neurons in the culture and this likely diluted our ability to detect a robust change in the quantified samples. Nonetheless, our data indicate different neuronal activity, including synaptic stimulation, increases Imp-β1 local translation in the synapto-dendritic compartment.

### Importin-β1 association with cargo proteins at the synapse

The activity-dependent local translation of Imp-β1 suggests the cargo binding of synaptic proteins to the nuclear import machinery is highly regulated. While activity-dependent movement of proteins have been previously described for several synapto-dendritic proteins, only a subset of these proteins are known to associate with nuclear adaptor complexes for its nuclear entry (Behnisch et al., 2011, Lai et al., 2008, Mikenberg et al., 2007, Dinamarca et al., 2016, Lim et al., 2017, Herbst and Martin, 2017).

To confirm if activity promotes Imp-β1 complex formation at the synapse, we performed co-immunoprecipitation (co-IP) experiments on synaptosomes using Imp-β1 as bait (**Fig. 6A**). This assay allows the identification of additional nuclear adaptor proteins or synaptic proteins that respond to neuronal activity by binding to Imp-β1. We purified forebrain synaptosomes from adult male mice and profiled the fractions via immunoblotting to identify fractions enriched with pre- and post-synaptic markers. As expected, the synaptosome fraction is enriched in both PSD95 and synaptophysin (SYPH) with the presence of Imp-β1 (**Fig. S6A**), all the mouse Imp-α isoforms (α1, α2, α3, α4 and α6) and importin-7 (IPO7) (**Fig. 6B-i**). Importantly, since we are looking for nuclear proteins at the synapse, no Histone-H3 was detected in the synaptosome fraction (**Fig. S6A**). Next, these synaptosomes were stimulated with KCl and glycine, a protocol known to trigger AMPA receptor insertion and local translation (Corera et al., 2009). Here, we show that the stimulation paradigm triggered calcium-dependent dephosphorylation of CRTC1 in stimulated synaptosomes (**Fig. S6B**). We performed co-IP on basal and stimulated synaptosomes and noted that IPO7 and all Imp-α isoforms (α1-4, α6), except for Imp-α2, associated with Imp-β1. The co-IP data suggests that the heterotrimeric interaction of Imp-α/β1-cargo is stable during affinity purification since the importin-beta binding (IBB) domain of Imp-α is inaccessible to Imp-β1 in the absence of cargo binding (Lott and Cingolani, 2011, Oka and Yoneda, 2018). Not surprisingly, we also observed an increased Imp-β1 protein level in stimulated synaptosomes and enhanced interaction between Imp-β1 and most of its binding partners (α1, α4 and α6, IPO7; **Fig. 6B-i**), likely reflecting an increased association of these nuclear adaptor proteins with Imp-β1 following activity-induced Imp-β1 local translation.

**Figure 6.**
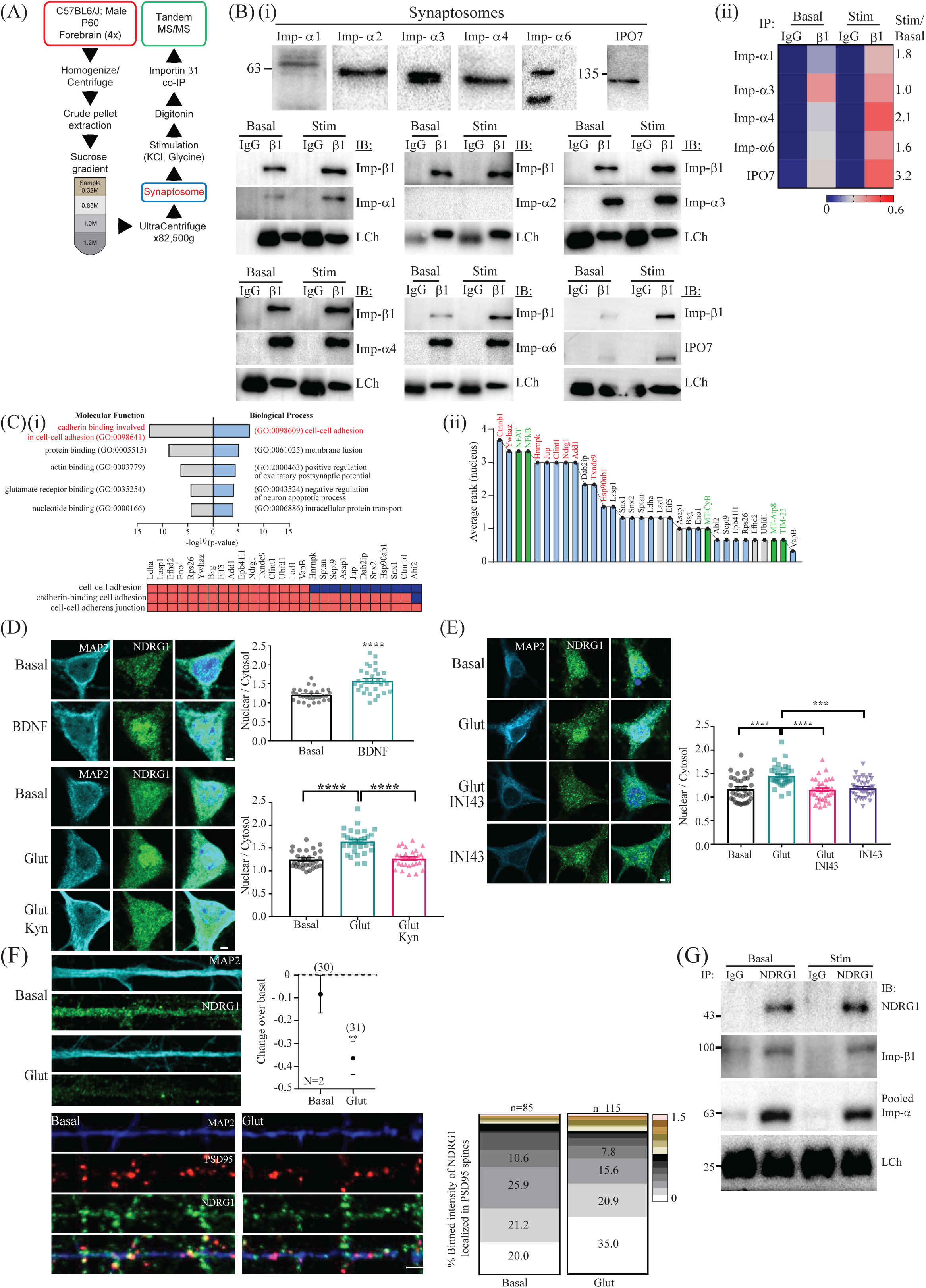
Imp-β1 association with NDRG1 at the synapse and undergoes Imp-β1 mediated nuclear accumulation following glutamatergic stimulation. **(A)** A schematic diagram showing the workflow of synaptosomal purification, stimulation, and Imp-β1 co-IP for tandem mass spectrometry (MS/MS) sequencing. **(B-i)** Lysates from purified synaptosomes (*upper panels*) and from Imp-β1 co-IPs (*lower panels*) were probed with antibodies targeted against Imp-α isoforms, IPO7 and IgG light chain (LCh). **(B-ii)** A heatmap showing average emPAI scores of Imp-α isoforms and IPO7 from three independent MS/MS sequencing data of Imp-β1 co-IP of basal and stimulated synaptosomes. (N=3) **(C-i)** GO analyses using DAVID showing the top enriched GO terms for molecular functions (MF) and biological processes (BP) from the candidate list (277 protein). The top ranked GO terms for each category are plotted against their *p*-values and listed in red. Candidate proteins from functional annotation clustering in DAVID associated with cell adhesion are listed alongside their individual GO terms. (**C-ii**) Proteins in (**C-i)** are scored and ranked for their nuclear presence using Compartments. Proteins with blue bars indicate synaptic presence while red fonts indicate nuclear presence. Green bars are control proteins (NFAT and Nf-κβ as positive controls for nuclear presence while MT-CyB, MT-Atp8 and TIM-23 as negative controls for mitochondrial-localized proteins). **(D)** Cultured mouse hippocampal neurons stimulated with BDNF (25 ng/mL, 1 h) as well as glutamate (Glut; 40 μM, 5 min) in the presence or absence of kynurenic acid (Kyn; 1 mM, 5 min) before fixing and immunolabeling with antibodies targeted against MAP2 (cyan), NDRG1 (green) and Hoechst nuclear dye (blue in merged image). The nuclear-to-cytoplasmic ratio of NDRG1 were quantified and the group data from independent experiments were plotted. Statistical analyses performed on group data use one way-ANOVA with Tukey’s post-hoc analyses for more than two experimental groups or unpaired T-test for two experimental groups (N=2; \*\*\*\**p* < 0.0001). Scale bar, 2 µm. **(E)** Cultured hippocampal neurons stimulated with glutamate (Glut; 40 μM, 5 min), in the presence or absence of INI43 (5 µM, 1 h), before fixing and immunolabeling with antibodies targeted against MAP2 (cyan), NDRG1 (green) and Hoechst nuclear dye (blue in merged image). The nuclear-to-cytoplasmic ratio of NDRG1 were quantified and the group data from independent experiments were plotted. Statistical analyses performed on group data use one way-ANOVA with Dunn’s post-hoc analyses (N=2; \*\*\**p* < 0.001, \*\*\*\**p* < 0.0001). Scale bar, 2 µm. **(F)** Cultured mouse hippocampal neurons stimulated with glutamate (Glut; 40 μM, 5 min) before fixing and immunolabeling with antibodies targeted against MAP2 (cyan or blue), PSD95 (red) and NDRG1 (green). *Upper panels*: The fold change of dendritic NDRG1 of stimulated against basal neurons were quantified and the group data from independent experiments were plotted (bracketed numbers indicate n number). Unpaired t-test analysis was conducted between basal and stimulated groups (N=2; \*\**p* < 0.01). *Lower panels:* The intensity of NDRG1 that colocalized with PSD95 (synaptic NDRG1) were quantified and binned. The percentage of each binned intensity category in basal and stimulated groups was quantified (N=2; Basal: n=85, Stimulated: n=115). Scale bar, 2 µm. **(G)** Co-IP using NDRG1 antibodies were performed in basal and stimulated synaptosomes and probed with antibodies targeted against NDRG1, Imp-β1, Imp-α isoforms and light chain (LCh).

To complement the immunoblotting data from co-IP experiments, we performed tandem mass spectrometry (MS/MS) sequencing on Imp-β1 co-IP samples, that were independent from the immunoblotting samples, and used exponentially modified protein abundance index (emPAI) to quantify the abundance of proteins in the samples. As expected, Imp-β1 protein is one of the highest enriched proteins for each of these co-IP experiments, with more Imp-β1 signal detected in the stimulated (average 2-fold increment) over the basal synaptosome samples. Consistent with the immunoblot data, the MS/MS results confirmed a robust Imp-β1 association with Imp-αs and IPO7 (**Fig. 6B-ii**). Specifically, Imp-β1 interactions with Imp-α1, α3, α4, α6 and IPO7 were readily detected in synaptosomes and all but Imp-α3 had a stimulation score of >1.0, with Imp-β1-IPO7 interaction showing the highest increase at 3.2-fold (**Fig. 6B-ii**). Importantly, the MS/MS emPAI scores for IgG co-IP controls detected negligible binding for all Imp-α isoforms and IPO7, indicating the high specificity in the identification of the protein complexes. Curiously, both immunoblotting and MS/MS sequencing data confirmed the absence of Imp-α2/β1 complex in either basal or stimulated synaptosomes (**Fig. 6B**). This is surprising and hints at the possibility of selective Imp-β1 heterodimerization that is stimulus or compartment specific.

We next analysed the MS/MS data for Imp-β1 co-IP to identify synaptic proteins that bound to Imp-β1 complexes. We ran triplicate co-IP samples in MS/MS and applied a series of stringent filters to obtain strong candidates for Imp-β1 binding. We considered only candidate proteins from the co-IP that registered no background binding to IgG controls (emPAI scores of -1). This criterion greatly reduces false positives even though some *bona fide* candidate proteins that bind to Imp-β1 may be excluded. Next, we shortlisted 277 candidate proteins which showed stable (Imp-β1 emPAI in stimulated sample > 0.1) and enhanced binding to Imp-β1 in stimulated synaptosomes (Imp-β1 emPAI stimulated > basal).

We then analysed the shortlisted candidate proteins using different methodologies. In the first approach, we compared the 277 candidate proteins against published databases. For synaptic presence, we cross-referenced the candidates against three databases: SynGO (Koopmans et al., 2019), SynaptomeDB (Pirooznia et al., 2012) and a published synaptosome proteomic database (Filiou *et al.* 2010). Synaptic proteins from all three databases share a significant overlap with 71 % (199/277) of our candidate proteins appearing at least once out of the three databases. We then compared the 277 candidates against the mouse nuclear proteome listed in MGI database and identified approximately 38 % (105/277) of our proteins annotated as nuclear proteins (**Fig. S6C-i**). Finally, we compared the two lists and isolated 83 proteins that shared dual localization in both the synapse and the nucleus (**Fig. S6C-i**), including members of the importin family.

Of the 83 proteins, more than one-third (34/83) are reported to undergo stimulus-driven nucleocytoplasmic shuttling, with several known to associate with IPO7 or the Imp-α/β1 import pathway, suggesting the possibility that some of these proteins may also respond to neuronal activity (**Fig. S7**). Indeed, three identified proteins, AIDA (Jordan et al., 2007), β-catenin (Chen et al., 2006, Abe and Takeichi, 2007, Schmeisser et al., 2009b), and ERK1 (Zhai et al., 2013, Wiegert et al., 2007) have been reported to shuttle into the nucleus from synapto-dendritic compartment in response to neuronal activity. Furthermore, a subset of these proteins are known to undergo activity-dependent nuclear shuttling in neurons, such as CDK5 (Liang et al., 2015) and Ywhaz (Neasta et al., 2012). In particular, Ywhaz is shown to interact with Imp-α2 and Imp-α4 (Arjomand et al., 2014). Other proteins such as Adducin-1 (Vukojevic et al., 2012), CASK (Malik and Hodge, 2014, Gillespie and Hodge, 2013), FMRP (Kanellopoulos et al., 2012, Tian et al., 2017), GSK-3β (Takashima, 2012), and HNRP K (Folci et al., 2014, Leal et al., 2017) are well-characterized plasticity proteins with dual functions in both the nucleus and the synapses.

In our second approach, we performed GO analyses on the candidate protein list (277 proteins) using DAVID (Huang da et al., 2009a, Huang da et al., 2009b) and the top GO terms for both molecular function and biological processes belonged to cell adhesion and cell junction proteins (**Fig. 6C & S6C-ii**). Functional annotation clustering show GO terms representing cell adhesion, cell adherens junction and cadherin-associated with cell binding enriched among the candidate proteins (**Fig. 6C**). We were intrigued that many cell adhesion and cytoskeleton-associated proteins bound to Imp-β1 at the synapse. Indeed, it is documented that this category of proteins in neurons undergo active nuclear import (Grabrucker et al., 2014, Birbach et al., 2006, Murk et al., 2012). We analysed the list of cell adhesion candidate proteins defined by DAVID using Compartments (Binder et al., 2014), a subcellular localization database, which generates an average rank for nuclear-localized proteins based on a combination of experimental evidence, text mining and software prediction. Proteins with a high average rank have a much higher likelihood to be localized in the nucleus. Nuclear proteins that bind to Imp-α/β1 such as NFAT and NF-κβ were included as positive controls and they have a high average rank (≥3) while mitochondrial proteins such as MT-CyB, MT-Atp8 and TIM-23 were used as negative controls with a much lower rank (≤1; both controls are shown in **Fig. 6C** with green bars). Reassuringly, many of the cell adhesion proteins in our candidate list with a high average rank also localized in the synapse (**Fig. 6C**; red fonts = annotated in nucleus; blue bars = annotated in synapse). Among the list of cell adhesion proteins, coupled with the information gathered in our first approach, we selected NDRG1 for further analysis.

### NDRG1 undergoes importin-β1 mediated nuclear accumulation following neuronal stimulation

The N-myc downstream regulated gene1 (NDRG1) is a synaptic protein identified in all three databases we cross-referenced and it belongs to the hydrolase superfamily (Schonkeren et al., 2019). More importantly, NDRG1 has been reported to undergo nucleocytoplasmic shuttling (Kitowska and Pawelczyk, 2010, Shi et al., 2013) and overexpression or knock-out studies of NDRG1 indicate the protein impacts gene expression (Schonkeren et al., 2019, Kovacevic et al., 2013, Cai et al., 2017). However, whether nuclear entry of NDRG1 directly modulates transcription in neurons is not known. In the peripheral nervous system, a nonsense mutation in NDRG1 is shown to be the underlying cause for peripheral neuropathy and Charcot-Marie-Tooth disease (Okuda et al., 2008). In the central nervous system, NDRG1 is found in developing hippocampal neurons as well as mature astrocytes in adult mice (Okuda et al., 2008), but its function is undefined.

Using antibodies validated against isoform-specific NDRG knockout mice (Okuda et al., 2008), we show NDRG1 expression in cultured hippocampal neurons (DIV14) and is localized in the somato-dendritic compartment with a mild nuclear enrichment under basal condition (**Fig. 6D**). Incubation with leptomycin B (LMB), which blocks *crm-1* nuclear export, led to a nuclear enrichment of NDRG1, verifying its nucleocytoplasmic shuttling property in neurons (**Fig. S6D**). We next stimulated cultured neurons with BDNF (25 ng/mL; 1 h) and observed a marked increase in nuclear NDRG1 (**Fig. 6D**). Similarly, stimulation with glutamate (40 µM, 5 min) also resulted in a robust nuclear enrichment of NDRG1, which is abolished in the presence of kynurenic acid, a broad-spectrum inhibitor of glutamatergic receptors (**Fig. 6D**). Crucially, the nuclear accumulation of NDRG1 is mediated by Imp-β1 as the addition of INI43 prior to and during glutamatergic stimulation abolishes the activity-induced nuclear accumulation of NDRG1 (**Fig. 6E**).

We next analyzed the NDRG1 localized in synapto-dendritic compartment and showed that in addition to the nuclear accumulation of NDRG1 upon glutamatergic stimulation, there is a corresponding reduction in the dendritic localized NDRG1 (**Fig. 6F**). We also measured the NDRG1 intensity colocalized to PSD95 positive puncta in basal and glutamate-stimulated neurons and demonstrate a loss of NDRG1 in the post-synaptic compartment (**Fig. 6F**). Notably, the proportion of PSD95 positive puncta without NDRG1 signal increased upon glutamatergic stimulation and accounted for 35 % of the total quantified puncta. The cumulative frequency plot also shows basal synapses having a much stronger NDRG1 signal as compared to the synapses in the glutamatergic stimulated neurons (**Fig. 6F**). Lastly, to validate the presence of NDRG1 at the synapse and its association with Imp-β1, we performed co-IP experiments using NDRG1 antibody on both the basal and the stimulated synaptosomes. We detected the interaction between NDRG1 and the Imp-α/β1 nuclear adaptor complex in the synaptosomes (**Fig. 6G**). In summary, our proteomic study identified a collection of synaptic and nuclear proteins that bound to Imp-β1 upon stimulation. A subset of these synaptic proteins has cell adhesion properties, of which NDRG1 is shown to directly interact with Imp-β1 to undergo activity-dependent nuclear import following glutamatergic stimulation.

## DISCUSSION

Our results confirm that activity-dependent movement of soluble proteins through the association with Imp-β1 is critical for transcription underlying long-term plasticity. Despite its implied involvement, the role of Imp-β1 in nuclear import during transcription-dependent plasticity has never been directly tested through loss-of-function studies. This is in part because a homozygous deletion of Imp-β1 in mice is embryonic lethal (Miura et al., 2006). To circumvent any long-term cellular dysfunction due to manipulations of Imp-β1 protein expression, we employed INI43, a small molecule inhibitor to block Imp-β1 nuclear entry just before and during neuronal stimulation (van der Watt et al., 2016). Our results on INI43 inhibiting Imp-β1 nuclear entry in neurons is consistent with other published studies and this inhibition is selective for Imp-β1 mediated nuclear import as activity-dependent entry of NFATc3, but not CRTC1, is impacted (Ch’ng et al., 2015, Wild et al., 2019). Moreover, in line with our hypothesis that Imp-β1 plays an important role in neuronal plasticity, the disruption of Imp-β1 function impaired multiple cellular processes associated with long-term plasticity, including CREB phosphorylation, JunB nuclear accumulation, induction of canonical activity-regulated genes expression and the maintenance of late-LTP.

Our experiments also indicate that Imp-β1 responds to synapse specific stimuli associated with LTD by entering the nucleus, opening up the possibility that LTD also requires the nuclear transport of soluble proteins to initiate transcription. In support of this idea, several reports have shown that LTD is transcription-dependent (Kauderer and Kandel, 2000, Kemp et al., 2013, O’Riordan et al., 2006), with certain forms of LTD, such as DHPG-mediated and mGluR-LTD, driving transcription of activity-regulated genes (Lindecke et al., 2006) as well as the activation of Nf-kβ nuclear signaling and DNA binding (O’Riordan et al., 2006, Wang and Zhuo, 2012). Furthermore, in other regions of the brain such as cerebellar Purkinje cells, long-term synaptic depression is transcription-dependent and requires the activation of MEF2 (Andzelm et al., 2019, Napolitano et al., 1999). In our hippocampal slice recordings, we observed a failure in inducing LTD when Imp-β1 function is blocked. This impairment is not due to the loss of synaptic transmission since INI43 is well-tolerated and does not alter baseline synaptic transmission even after 180 min post-stimulation. Hence, the likely possibility is that the loss of Imp-β1 function impairs the signaling mechanism required for the induction or maintenance of LTD. For example, it is known that PP2A is critical for LTD induction through the dephosphorylation of the GluA1 subunit of the AMPA receptor (Mauna et al., 2011). Neurons express an endogenous PP2A inhibitor (I_2_^PP2A^) which encodes a canonical NLS and is basally sequestered in the nucleus. A loss of Imp-β1 function would result in the cytoplasmic retention of I_2_^PP2A^ and the subsequent inhibition of PP2A activity on AMPA receptors required for LTD induction (Arif et al., 2014). Alternatively, the pre-existing association of target synaptic proteins with the nuclear adaptor proteins including Imp-β1 may be crucial for the induction of LTD. Indeed, proteins from our target list that associated with Imp-β1, such as GSK3β, Camk2α or MAPK, are all required for the induction of LTD and small molecule inhibition of these plasticity proteins are shown to block LTD induction and abolish late-LTD (Sajikumar et al., 2007, Peineau et al., 2007). It is also possible that Imp-β1 may have other local but uncharacterized role during long-lasting synaptic plasticity including reverse phase separation via disaggregation of NLS-bearing proteins or as potential interactors during LTD-induced synaptic tagging (Guo et al., 2018).

While our data supports activity-dependent entry of soluble proteins in the nucleus, it is not clear which Imp-β1 heterodimer(s) is responsible for this role. In this regard, Imp-β1 is unique as it bridges both the classical and non-classical nuclear import pathways and there is evidence that both pathways are involved in maintaining long-term plasticity. We and others have shown that the classical Imp-α/β1 complex responds to activity by entering the nucleus and regulating neuronal plasticity (Jeffrey et al., 2009, Thompson et al., 2004). Meanwhile, IPO7, a member of the Imp-β superfamily, is reported to shuttle phosphorylated ERK (pERK) into the nucleus in an activity-dependent manner (Li et al., 2016, Chuderland et al., 2008). Of note, IPO7 heterodimerization with Imp-β1 may be critical for the nuclear entry of pERK (Lorenzen et al., 2001) and our data show enhanced heterodimerization of IPO7-Imp-β1 and association with ERK1 in stimulated synaptosomes (**Fig. 6B-ii & S7**).

Establishing the importance of activity-driven Imp-β1 nuclear import during long-term plasticity provides a framework for us to study the contribution of Imp-β1 nuclear signalling from different subcellular compartments, particularly at the synapto-dendritic regions where an increasing number of synapto-dendritic proteins reported to undergo activity-dependent retrograde movement to the nucleus (Herbst and Martin, 2017, Lim et al., 2017, Marcello et al., 2018, Parra-Damas and Saura, 2019, Brigidi et al., 2019). Since many signalling cascades at the synapto-dendritic compartment are activity-dependent and highly regulated, we reasoned that cargo binding by importins is likely to be regulated by local mechanisms. In the paper, we provide evidence that Imp-β1 function at this subcellular compartment is regulated by local translation. We report the presence of at least three distinct Imp-β1 mRNA isoforms in the synaptosomes and using different assays including Puro-PLA in the presence of CHX, we observed activity-dependent increase in Imp-β1-ribosome complex in the synapto-dendritic compartment (David et al., 2012, tom Dieck et al., 2015). While we hypothesized that the increase in Imp-β1-ribosome signal could be asymmetrically distributed at synapses (Group IV puncta; **Fig. 5D-ii**) during bicuculline treatment, we nonetheless observed an equal distribution in the increase of signal in all the synapto-dendritic compartments with a trend toward synaptic localization. We reason that the heterogeneity in bicuculline response in the hippocampal cultures prevented a more robust quantification of the Imp-β1-ribosome shift in its synaptic distribution.

Collectively, our data, together with the published report by Jeffrey and colleagues (2009), argue that importin function at the synapse is highly regulated and likely shaped by activity. Our updated model predicts that synaptic stimulation triggers the activation of NMDA receptors and phosphorylation of the NLS in the NR1a cytoplasmic tail by protein kinase C which in turns, releases Imp-α1 anchored at the site. In parallel to Imp-α1 release, Imp-β1 mRNA is loaded onto ribosomes and translated in the synapto-dendritic compartment. The overall increase in local importins concentration will enhance the formation of different Imp-β1 heterodimers, including those with Imp-α and IPO7, which then facilitate the binding to different synaptic proteins for retrograde movement and entry to the nucleus (**Fig. 7**). Interestingly, a recent report identifying translating mRNAs in the neuropils detected the presence of different importin mRNAs, including Imp-β1, suggesting that local translation of other importins, apart from Imp-β1, can occur at the synapse (Biever et al., 2020).

**Figure 7.**
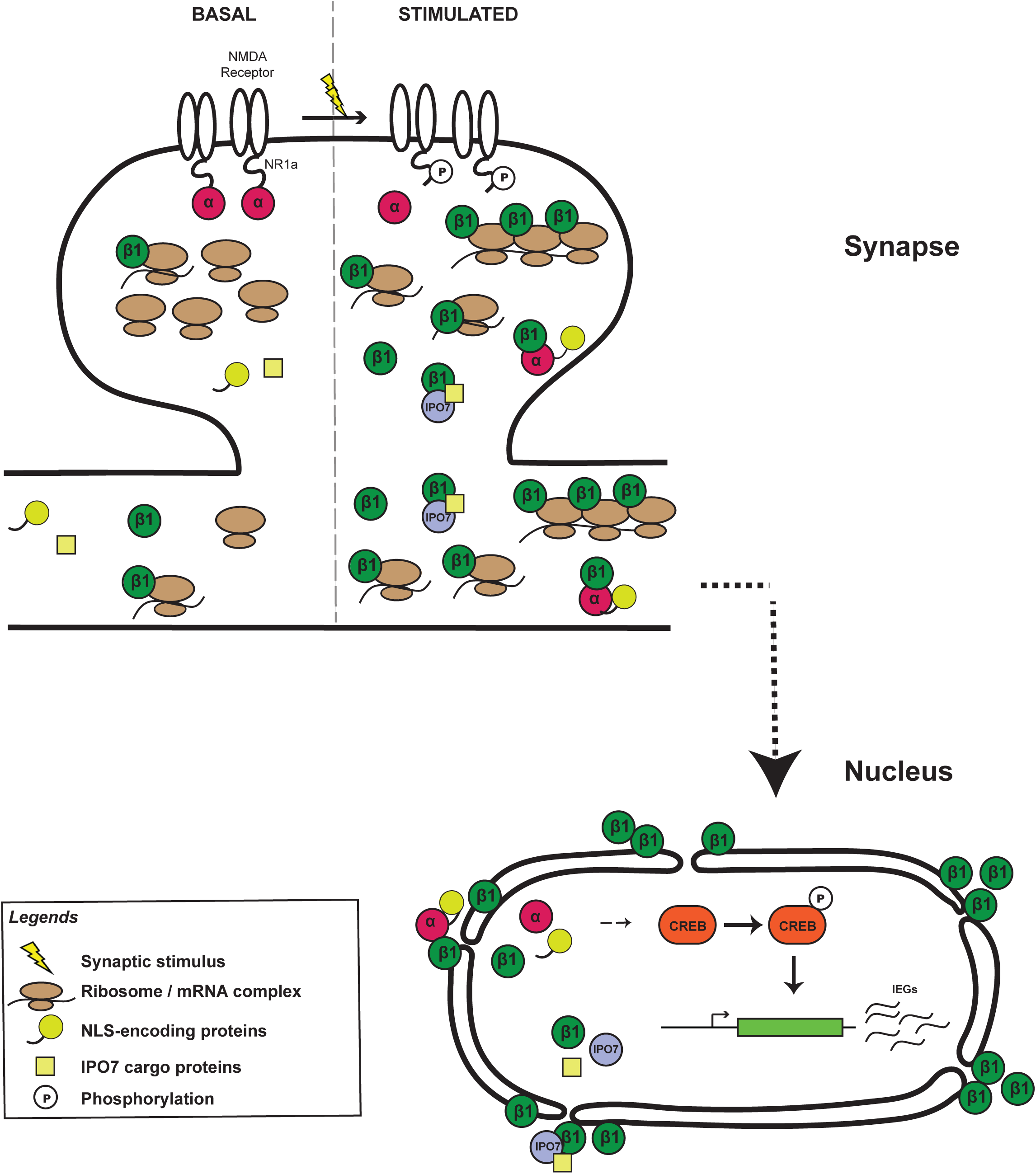
Model for activity dependent regulation of importins at the synapto-dendritic compartment. A model for the activity-dependent regulation of Imp-α isoforms and Imp-β1 at the synapto-dendritic compartment and the subsequent nuclear transport and entry of cargo-bearing Imp-β1 adaptor complexes to regulate transcription of activity-regulated genes.

In this study, we decided to cast a wide net to identify additional synaptic proteins that bound to Imp-β1 heterodimers in the stimulated synaptosomes. Our MS/MS sequencing and immunoblots of Imp-β1 co-IP in basal and stimulated synaptosomes showed Imp-β1 association with IPO7 and all Imp-αs except Imp-α2 (**Fig. 6B**). Since the Imp-β1 co-IP generated a diverse target list, stringent selection criteria were necessary to remove background noise. Among our list of candidate proteins, at least three proteins, β-catenin, AIDA and ERK1 are proteins previously identified in the synapto-dendritic compartment that undergo long distance movement to the nucleus in response to activity (Jordan et al., 2007, Zhai et al., 2013, Schmeisser et al., 2009a). Interestingly, functional annotation clustering of the target list showed enrichment for GO terms associated with cell-cell adhesion. Perhaps, this is not surprising given that the nucleocytoplasmic shuttling of cell adhesion proteins or cytoskeletal proteins have been well-documented in many cell types, including neurons. A subset of these proteins respond to different stimulation by undergoing nuclear translocation, such as Shank3, which is implicated in ASD (Grabrucker et al., 2014, Jiang and Ehlers, 2013), Profilin, which is involved in Fragile X syndrome (Birbach et al., 2006, Murk et al., 2012, Michaelsen-Preusse et al., 2016), Afadin, where its deletion leads to hydrocephalus (VanLeeuwen et al., 2014, Yamamoto et al., 2013) and Vimentin, implicating in Alzheimer’s disease (Perlson et al., 2005, Levin et al., 2009). Incredibly, many of these dual-localized proteins not only help maintain the structural integrity of different subcellular compartments, they also function as transcriptional modulators capable of regulating gene expression (Kumeta et al., 2012). We selected one of the synaptic cell adhesion proteins, NDRG1 for further studies and show that it exhibits dual localization in the synapse and the nucleus. Our characterization shows that NDRG1 undergoes Imp-β1-mediated nuclear accumulation during glutamatergic stimulation with a corresponding loss in the synapto-dendritic compartment. Further work will be performed to ascertained synapse-to-nucleus signaling of NDRG1.

## ACKNOWLEDGEMENTS

We thank Albert I. Chen, Kwok On Lai for proof-reading the manuscript, Koichi Kokame, Kelsey Martin, Mark Dell’Acqua, Yuh Min Chook for generously sharing advice and reagents, Balakrishnan Kannan for microscopy help and Jessica Ruth Gaunt for proteomic analyses.

## AUTHOR CONTRIBUTIONS

Conceptualisation, Y.J.L, T.H.C; Validation, Format analyses, Investigation, C.F.T, S.K.S (Mass spectrometry), S. N., S.S. (Slice electrophysiology), Y.J.L (others); Writing-original draft and Visualization, Y.J.L, T.H.C; Writing-review and edition, S.S, T.H.C, S.K.S; Supervision, T.H.C; Funding acquisition T.H.C.

## EXPERIMENTAL PROCEDURES

### Animals

All experiments were performed using protocols approved by Nanyang Technological University Institutional Animal Care and Use Committee (IACUC #E0019 and #A18091). C57BL/6 mice and Sprague-Dawley rats were used. Mouse and rat hippocampal neurons were isolated from P0 to P1 pups and maintained for 14 to 17 days in-vitro (DIV) prior to any experiments. Synaptosomes and acute slices were isolated from the forebrain of P56 to P60 male mice.

### Antibodies

| Antibodies | Source | Dilution |
| --- | --- | --- |
| Alexa Fluro conjugated secondary Ab | Life Technologies | 1:1000 (ICC) |
| Chicken-anti-MAP2 | Phosphosolutions | 1:10,000 (ICC) |
| HRP-linked anti-mouse and anti-rabbit secondary Ab | Cell Signaling | 1:1000 (IB) |
| Mouse specific light chain (HRP conjugated) | EMD Millipore | 1:1000 (IB) |
| Mouse-anti-GAPDH | ProteinTech | 1:5000 (IB) |
| Mouse-anti-HA | Sigma | 1:1000 (IB) |
| Mouse-anti-importin-β1 (3E9) | Thermo Fisher | 1:1000 (ICC); 3μg (IP); 1:1000 of PLA probe conjugated 3E9 |
|  |  | (420 µg/ mL) |
| Mouse-anti-Imp-α1 | Santa Cruz | 1:200 (IB) |
| Mouse-anti-IPO7 | Santa Cruz | 1:200 (IB) |
| Mouse-anti-JUNB | Santa Cruz | 1:100 (ICC) |
| Mouse-anti-PSD95 | Invitrogen | 1:5000 (IB);<br>1:1000 (ICC) |
| Mouse-anti-puromycin | Kerafast | 1:1000 of PLA<br>probe conjugated<br>antibody (420 µg/<br>mL) |
| Mouse-anti-synaptophysin<br>(Clone SY38) | EMD Millipore | 1:1000 (IB) |
| Mouse IgG | Sigma | 3µg (IP) |
| Rabbit-anti- Imp-α2-4, 6 | Gift from Martin K.C | 1:10,000 |
| Rabbit-anti-cleaved caspase 3 | Cell Signaling | 1:400 (ICC) |
| Rabbit-anti-CRTC1 | Bethyl labs | 1:3000 (ICC) |
| Rabbit-anti-GFP | Thermo Fisher | 1:1000 (ICC) |
| Rabbit-anti-H3 | Cell Signaling | 1:1000 (IB) |
| Rabbit-anti-NDRG | Gift from Kokame, K | 1:100 (ICC); 3µg<br>(IP) |
| Rabbit-anti-pCREB (S133) | Cell Signaling | 1:500 (ICC) |
| Rabbit-anti-synaptophysin | EMD Millipore | 1:500 (ICC) |
| VeriBlot for IP Detection<br>Reagent (A131366) | Abcam | 1:1000 (IB) |

### Plasmids and viruses

FLAG-tagged Imp-α (1-4,6) plasmids. HA-tagged human Imp-β1 plasmid was a kind gift from Chook, Y. M (UT Southwestern) and used to generate Imp-β1-Dendra2 (L22-Camk2α-hImp-β1 and L22 L22:Camk2a-GFP; GenScript BioTech, Hong Kong). sGFP-hNFATc3 plasmid was a kind gift from Mark Dell’Acqua (U. of Colorado, Denver). pFCK(1.3)GW was a kind gift from Pavel Osten (Addgene plasmid #27230; http://n2t.net/addgene:27230).

### Mammalian cells and hippocampal neuronal cultures

Hippocampal neurons were isolated as previously described in (Ch’ng et al., 2012) and plated on poly-DL-lysine (0.5 mg/ml; Sigma) coated coverslips (Carolina Biologicals, Burlington, NC), maintained in Neurobasal media, (Invitrogen, Carlsbad, CA), supplemented with B27 (Invitrogen), GlutaMAX (Invitrogen), fetal bovine serum (FBS) and gentamycin (Life Technologies) one day post-culture for mouse cultures. For rat cultures, Neurobasal media was supplemented with B27, GlutaMAX, monosodium glutamate (Sigma) and β-mercaptoethanol (Sigma) during plating and maintained in Neurobasal media supplemented with B27 and GlutaMAX for the remainder of the culture (Ch’ng et al., 2012). For most experiments, unless otherwise indicated, neurons were incubated with various pharmacological agents in the conditioned neuronal media, kept in humidified incubator (37°C, 5% CO_2_) for the appropriate amount of time before cells were processed for immunoblotting or immunocytochemistry. HEK293T cells were grown in Dulbecco’s Modified Eagle Media (DMEM; Nacalai) supplemented with 10% fetal bovine serum (FBS; Hyclone) and 1% Penicillin/Streptomycin mixed solution (Nacalai), kept in a humidified incubator (37°C, 5% CO_2_) for the appropriate amount of time before experiments

### Pharmacological treatment

The following pharmacological agents were used: APV (40 μM; Tocris), Bicuculline (Bic, 40 or 50 μM; Sigma), Brain-derived neurotrophic factor (BDNF, 50 ng/mL), Cycloheximide (CHX, 50 µM; Sigma), Glutamate (40 μM; Sigma), Glycine (20 or 100 μM; Sigma), H_2_O_2_ (500 µM, Sigma), INI-43 (5 μM; Sigma), Kynurenic acid (Kyn, 1 mM, Sigma), Leptomycin B (LMB, 20 nM; Sigma), MG132 (5 µM), N-ethylmaleimide (NEM, 50 µM; Sigma), N-Methyl-d-aspartic acid (NMDA, 20 µM), Potassium Chloride (KCl, 50 mM; Merck), (RS)-3,5-Dihydroxyphenylglycine (DHPG, 100 µM; Tocris), Tetrodotoxin (TTX, 1 μM; Tocris), 4-Aminopyridine (4AP, 200 μM). Inhibitors were added to neuronal cultures prior to stimulation as described in each experiment. For experiments involving KCl stimulation, hippocampal culture medium was replaced either with basal artificial cerebrospinal fluid (aCSF; 126 mM NaCl; 24 mM NaHCO_3_; 1 mM NaH_2_PO_4_; 2.5 mM KCl; 2 mM CaCl_2_; 2 mM MgCl_2_; 10 mM D-Glucose; 0.4 mM ascorbic acid) or high K^+^ no Mg^2+^ aCSF (78.5 mM NaCl; 24 mM NaHCO_3_; 1 mM NaH_2_PO_4_; 50 mM KCl; 10 mM CaCl_2_; 0 mM MgCl_2_; 10 mM D-Glucose; 0.4 mM ascorbic acid) during pharmacological treatments. For experiments involving the addition of NEM, hippocampal culture medium was replaced with Tyrode’s solution during pharmacological treatments (140 mM NaCl, 10 mM HEPES, 5 mM KCl, 3 mM CaCl_2_, 1 mM MgCl_2_, 10 mM glucose).

### Forebrain Acute Slices

Mouse forebrain coronal acute slices (250 μm) were sectioned using vibratome in cold cutting solution, gassed with 5 % CO_2_ and 95 % O_2_. The slices were then allowed to recover for three hours, in aCSF (126 mM NaCl, 24 mM NaHCO_3_, 1 mM NaH_2_PO_4_, 10 mM D-Glucose, 2.5 mM KCl, 2 mM CaCl_2_, 2 mM MgCl_2_, and 0.4 mM ascorbic acid) at 32°C, at 5 % CO_2_ and 95 % O_2._ Chemical induction of LTP in aCSF lacking MgCl_2_ (78.5 mM NaCl; 24 mM NaHCO_3_; 1 mM NaH_2_PO_4_; 50 mM KCl; 10 mM CaCl_2_; 0 mM MgCl_2_; 10 mM D-Glucose; 0.4 mM ascorbic acid; 40 µM forskolin) for 15 min, followed by high KCl (50 mM) for 10 min (Ch’ng et al., 2012). All slices were then used for immunocytochemistry experiments.

### Cellular transfection and transduction

Calcium phosphate transfection of cultured hippocampal neurons (DIV13-16) were conducted as previously published (Dudek et al., 1998). In brief, cultured neurons in 24-wells plate were incubated for 20-60 min with the DNA/calcium phosphate precipitate in neurobasal media before returned to conditioned media. Unless otherwise stated, transfected constructs neurons are allowed 16-24 h of expression prior to experiments. Transfection of plasmids in HEK 293T cells was carried out with DNAfectin™ (Invitrogen) following manufacturer’s protocols. Length of expression of these constructs will vary depending on the experiments and will be individually described. For lentiviral transduction of cultured hippocampal neurons, the viral inoculum was diluted in 250 µl of conditioned medium and incubated with the neurons (DIV 11) overnight, before swapping into the remaining conditioned medium without viral inoculum. In general, lentiviral integration and expression were allowed for a minimum of 3 days post-transduction before conducting any further experiments.

### Puromycin Proximity Ligation Assay (Puro-PLA)

Puro-PLA on cultured hippocampal neurons were conducted according to previously published protocols (tom Dieck et al., 2015). Briefly, after stimulation. all samples were first treated with cycloheximide (CHX; 355 µM, 30 min) to terminate nascent chain elongation and to prevent release of nascent peptide from ribosome (David et al., 2012, tom Dieck et al., 2015). This is followed by puromycin (3 μM, 15 min) treatment in conditioned medium in a humidified incubator (37 °C, 5 % CO_2_). Incubation was stopped by immediate fixation with 3.2 % PFA diluted in Tyrode’s solution. After fixation, the neurons were washed, permeabilized and treated for PLA following protocols described in the Duolink^®^ PLA Fluorescence kit (Sigma). Both the antibodies targeted against Imp-β1 and puromycin were conjugated directly to the PLA probes as PLA^plus^ and PLA^minus^ respectively using Duolink™ In Situ Probemaker PLUS and MINUS kits (Sigma). Neurons were then fixed with 3.2 % PFA before processing with immunocytochemistry.

### Synaptosome extraction and stimulation

Synaptosomes were isolated from for male mice in each extraction as described in Jeffrey *et al*. (2009) with minor modifications including: 18 up-and-down strokes for the first homogenization; 6 up-and-down strokes for the second homogenization; 700 x g first round centrifugation; crude pellet was homogenized in 4.5 mL of Solution B and layered on the discontinuous sucrose gradient and subsequently ultracentrifuged (82,500 x g for 2 h 15 min). Synaptosomes were harvested at the sucrose interface between 1.0 M and 1.2 M and 2.5 μM of CaCl_2_ were added to the purified synaptosomes before being divided for further stimulation. Activation of synaptosomes involves a 5 min incubation with 100 μM glycine followed by another 5 min incubation with 50 mM KCl. Cycloheximide (CHX, 50 µM) was added to the specified group 30 min before stimulation. The synaptosomes were kept in cold except during the stimulation. These synaptosomes were then used for qPCR analysis or immunoprecipitation experiments described below.

### Immunoprecipitation (IP)

Cold lysis buffer (0.025% digitonin, 150 mM NaCl, 10 mM Tris-HCl-pH 7.4), supplemented with protease inhibitor cocktail (Roche), was added to synaptosomes in a 1:1 ratio and rotated end-to-end in cold room for 30 min prior to IP assays. For HEK 293T cells, the lysates were prepared by scraping the (transfected) cells in PBS, followed by centrifugation at 5000 rpm for 3 min. Subsequently, the cold lysis buffer (1% NP-40, 50 mM Tris-HCl pH 8.0, 150 mM NaCl supplemented with protease inhibitor cocktail) was added to the cell pellet and incubated for 15 min on ice. Following lysis, 3 μg of the appropriate antibodies were added to each sample and incubated overnight, followed by 4 h end-to-end incubation with 300 μg /mL of Protein G magnetic beads (Thermofisher) in 4 °C. The beads were placed on a magnetic plate for 15 min on end and washed 3X with cold lysis buffer before resuspending the samples with 2X sample buffer and boiled at 105°C for 10 min before proceeding to immunoblotting or tandem mass spectrometry sequencing.

### Quantitative PCR (qPCR)

Total RNA was extracted with TRIzol (Invitrogen) and column purified with PicoPure RNA isolation kit (Cultured neurons; Thermofisher) or RNeasy Mini Kit (synaptosome and cytosolic fraction from homogenized mouse forebrain; Qiagen). For RT-PCR, cDNA was generated using RevertAid First strand cDNA synthesis kit (Thermofisher) with either poly-dT or random hexamer priming. SYBR Green Master Mix (Invitrogen) with specific primer pairs (table 1) were used for all qPCR assays and analyzed via QuantStudio 6 Flex Real-Time PCR System (Applied Biosystems). Technical and experimental triplicates were performed for all samples. For cultured neurons analyses, ΔCt values were calculated by normalizing the average raw qPCR results for each of the gene against neuron specific HPRT1. ΔΔCt values were then calculated by normalizing the ΔCt of the treated samples against basal controls. For synaptosomal and cytosolic Imp-β1 isoforms analyses, the mean Ct values obtained from primers targeted against long (L1 and L2) and full-length (F1 and F2) isoforms were normalized against the mean Ct values obtained from common primer (S1).

**Table 1:**
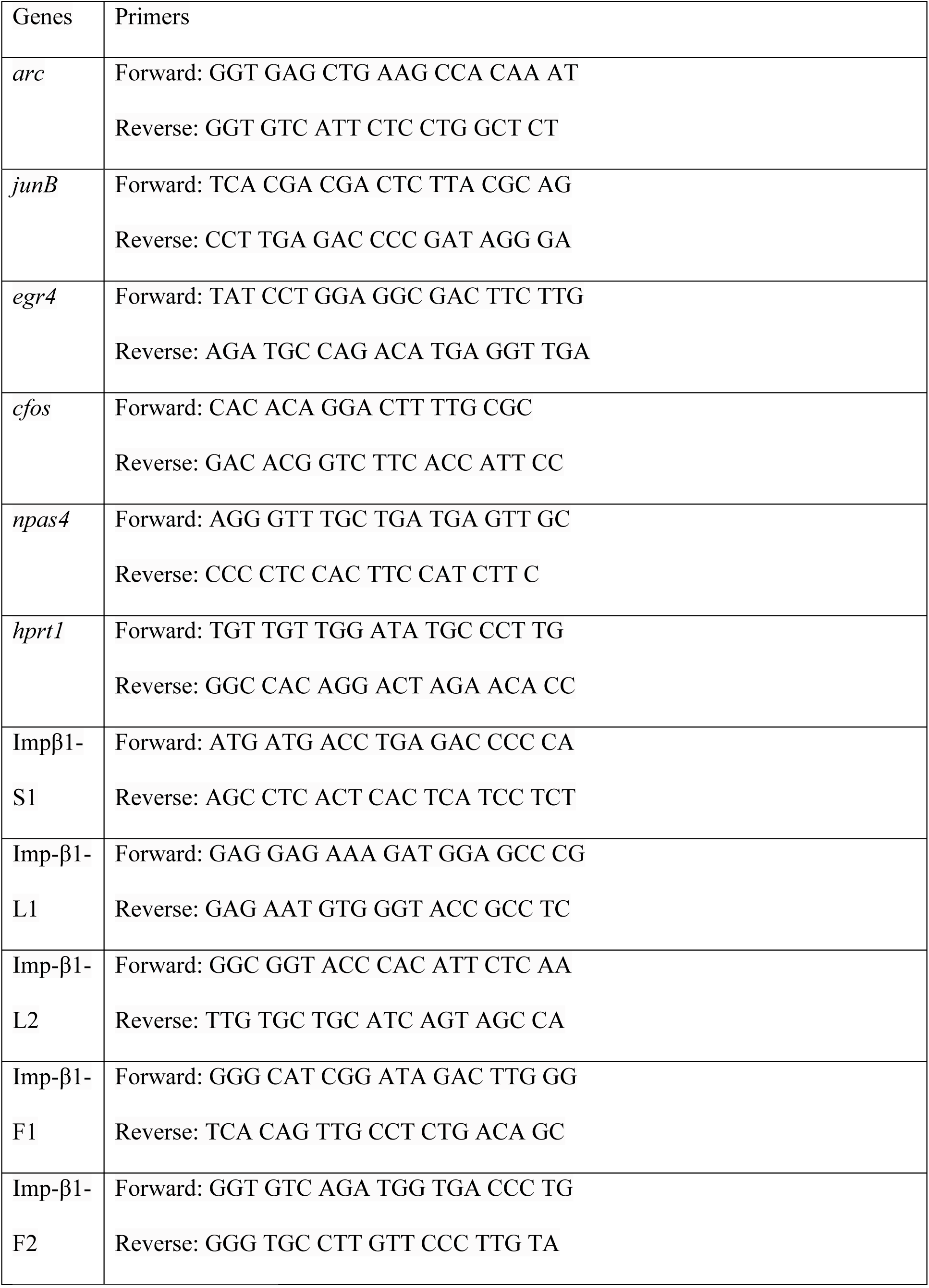
qPCR primer pairs.

### Immunoblotting (IB)

For HEK 293T cell lysates that were not prepared for co-IP experiments, HEK 293T cells were harvested in RIPA buffer (150 mM NaCl, 1 % TX-100, 0.5% sodium deoxycholate, 0.1 % sodium dodecyl sulfate, 50 mM Tris pH 8.0). Protein concentration was determined using Pierce BCA Protein Assay Kit (Thermofisher) according to manufacturer’s protocol where needed. All proteins were resolved in 10 % to 12 % SDS-PAGE and transferred to Immobilon transfer membrane (EMD Millipore). Membranes were blocked in ChemiBlocker (EMD Millipore) for at least 1 h before incubating in primary antibodies overnight at 4 °C followed by HRP-conjugated secondary antibodies for at least 1 h at room temperature. All membranes were washed three times with 0.1 % Tween 20 in PBS, 20 min/wash, after primary and secondary antibody incubation. SuperSignal™ West Pico PLUS Chemiluminescent Substrate (Thermo Scientific) was then added to all blots and imaged immediately with Bio-Rad imaging system with analyses and quantifications of bands performed using Image Lab™.

### Immunocytochemistry (ICC)

Cultured hippocampal neurons were fixed in 4 % paraformaldehyde diluted in Tyrode’s buffer (140 mM NaCl, 10 mM HEPES, 5 mM KCl, 3 mM CaCl_2_, 1 mM MgCl_2_, 10 mM glucose) and permeabilized in PBS containing 0.1% TX-100 and blocked in PBS in 10% goat serum. Mouse forebrain slices were fixed in 4% PFA diluted in aCSF (126 mM NaCl; 24 mM NaHCO_3_; 1 mM NaH_2_PO_4_; 2.5 mM KCl; 2 mM CaCl_2_; 2 mM MgCl_2_; 10 mM D-Glucose; 0.4 mM ascorbic acid) and permeabilized in PBS containing 0.4% TX-100 with blocking in PBS and 10% goat serum (in the presence of 0.2% TX-100). All primary antibody incubations were performed overnight in 4 °C and secondary antibodies staining were done 1 h at room temperature. For all preparations, samples were washed 3X with PBS between each step of the protocol. After antibody incubations, samples were mounted on slides using Aqua-Poly/Mount (PolySciences).

### Electrophysiology

Acute hippocampal slices were prepared from 5 to 7-week-old wild type BL-6 mice for electrophysiological recordings. Briefly, after anesthetization using CO_2_, the mice were decapitated, and the brains were quickly removed and transferred into cold, oxygenated aCSF. The aCSF contained the following (in millimolars): 124 NaCl, 3.7 KCl, 1.0 MgSO_4_·7H_2_O, 2.5 CaCl_2_·2H_2_O, 1.2 KH_2_PO_4_, 24.6 NaHCO_3_, and 10 D-glucose, equilibrated with 95% O_2_–5% CO_2_ (carbogen; total consumption 16 L h^−1^). From each hippocampus, 8–10 transverse hippocampal slices (400 μm thick) were prepared using a manual tissue slicer. The slices were incubated in an interface brain slice chamber (Scientific Systems Design, Mississauga, Ontario, Canada) at 32 °C for three to four hours at an aCSF flow rate of 1 mL min^−1^. For INI43 treated slices, aCSF containing 5 µM INI43 was bathed in the slices 30 min prior and 30 min post stimulation/depression.

For stimulation, a monopolar stainless-steel electrode (5 MΩ; A-M Systems, Sequim, WA, USA) was positioned within the stratum radiatum of the CA1 region. For the recordings of fEPSP (measured as its slope function), a monopolar stainless-steel electrode (5 MΩ; A-M Systems, Sequim, WA, USA) was placed in the CA1 apical dendritic layer. Signals were amplified by a differential amplifier (Model 1700; AM Systems) and digitized using a CED 1401 analog-to-digital converter (Cambridge Electronic Design, Cambridge, UK) and monitored online.

Input–output relation (afferent stimulation vs. fEPSP slope) was determined for each slice and the stimulus intensity that evokes 40% of the maximum fEPSP slope is set as test stimulus intensity. For baseline recording and testing at each time point, four 0.2-Hz biphasic constant-current pulses (0.1 ms per polarity) were used. L-LTP was induced using three stimulus trains of 100 pulses (‘strong’ tetanus [STET], 100 Hz; duration, 0.2 ms per polarity; intertrain interval, 10 min). L-LTD was induced using a strong low-frequency stimulation (SLFS) protocol of 900 bursts [one burst consisted of three stimuli at 20 Hz, and the interburst interval was 1 s (i.e., f = 1 Hz; stimulus duration, 0.2 ms/half wave; total number of stimuli, 2700)].

### Confocal imaging and analyses

Unless otherwise stated, all confocal images, either static or live cell, were performed on a Zeiss LSM800 (Carl Zeiss, Inc.) inverted scanning confocal microscope using either 40 X or 63 X 1.4 NA oil objectives. For live cell imaging of lysosomes, cultured mouse hippocampal neurons grown on glass coverslips were incubated in N-ethylmaleimide (NEM, 50 µM, 10 min) in Tyrode’s solution, followed by the addition of Lysotracker Deep Red (1 μM, 10 min, Thermofisher) before washout, and transferred to a live-cell imaging buffer (140 mM NaCl, 10 mM HEPES, 5 mM KCl, 3 mM CaCl_2,_ 0.1 mM MgCl_2,_ 10 mM glucose, 10 nM Ascorbic acid, pH 7.35). Cells were imaged on a heated stage assembly and images were acquired at 1s / frame for 60 s. The displacement length and cumulative displacement of the lysosomes was calculated using Imaris “track displacement length” plugin.

For live cell imaging of Imp-β1, rat hippocampal neurons were transfected with Imp-β1-Dendra2 constructs (DIV 13-16). After 16 h of expression, neurons were transferred to a buffer containing low (for basal) or no (for stimulation) Mg^2+^ Tyrode’s solution (140 mM NaCl; 10 mM HEPES pH 7.3; 5 mM KCl; 3 mM CaCl_2_; 0.1 mM MgCl_2_; 10 mM glucose; 10 nM Trolox pH 7.35) in the presence or absence of bicuculline (Bic; 40 µM). The entire assembly was placed on a heated stage for imaging. Neurons with comparable expression levels were selected for imaging. Three regions of interest (15 µm x 5 µm) on dendrites within 60 μm away from the soma were converted by brief exposure to UV laser three times (405 nm, 6.0%, scan speed: 3). Immediately after the photoconversion, photoconverted Dendra2 signal in the soma was tracked for 15 minutes (1 frame / min: t=1-15). The photoconverted signal from Dendra2 and Imp-β1-Dendra2 signal in the nucleus (based on pre-capture, low intensity snapshot of Hoechst) were quantified and normalized (t=1) across all time points in each condition.

For all confocal micrographs, quantifications were done using ImageJ unless otherwise stated. For nuclear-to-cytoplasmic ratio, cytoplasmic intensity was calculated by subtracting the nuclear mask (based on Hoechst) from the whole cell mask (based on MAP2). All dendritic measurements were obtained from dendrites taken 70 µm from the base of the cell body (unless otherwise stated). For the Puro-PLA dendritic assay, signals were processed with a constant threshold applied within the same experimental set using threshold function in ImageJ before quantifying for its integrated intensity.

Specifically, for Puro-PLA synaptic experiments, confocal images of dendrites taken 50 µm from the base of the cell body were captured in a fixed 50 µm x 15 µm field using 63X 1.4 NA oil DIC Plan Apochromat AiryScan objective lens on an LSM800 scanning confocal microscope, attached to a 32-element GaAsP-PMT array detector running the AiryScan module. Images were then deconvoluted using AiryScan deconvolution algorithm post-acquisition and the colocalization analyses were performed using IMARIS 6.3.1, with its surface-surface colocalization plugin. Prior to volumetric analysis, a constant threshold to remove background was used within each experimental set for each fluorescence channel: MAP2 (10% lower intensity threshold); PSD95 and synaptophysin (15-20% lower intensity threshold and split object); Imp-β1-Puro PLA (30% lower intensity threshold, smoothing and split object). Within each dendritic field, a 3-D volumetric surface was independently generated for all neuronal markers and PLA signals. The marker surfaces were then mapped onto the PLA surfaces to generate a 3-D mask that shows overlapping regions (**Fig. 4C**).

For analyses of Imp-β1 in spines (**Fig. 5B**), the spines were identified by thresholding the intensity and circularity of GFP puncta that are found within 1 µm away from the dendrites.

### Statistical analyses

All statistical analyses and graphs were done using GraphPad Prism 7.0. For two comparison groups, parametric t-test or Mann-Whitney non-parametric t-test were performed unless otherwise stated. For more than two comparison groups, one-Way ANOVA coupled with Tukey’s post-hoc analysis (parametric) or Dunn’s post-hoc analysis (non-parametric) was conducted unless otherwise specified. To test for normality, Shapiro-Wilk normality test was conducted for small sample size (n<30) while D’Agostino & Pearson normality test was conducted for large sample size prior to One-Way ANOVA test. Α probability value of α = 0.05 was used to determine statistical significance. All imaging and data collection are performed using appropriate cellular markers to select for specific cell types and to select viable cells without prior knowledge of experimental conditions. Unless otherwise stated, all data points are included in every analysis. Since data are obtained blind, specific data points are excluded only under extreme circumstances and only when they deviate significantly from the mean distribution (artifacts of staining, etc). In which case, an outlier test is performed with Prism GraphPad using ROUT method, Q = 1%. Specifically, an outlier test was conducted for NDRG1 dendritic analyses (**Fig. 6F**) and Imp-β1 localization in KCl stimulated neurons (**Fig. 1A**).

### Co-immunoprecipitation and LC-MS/MS analysis

Synaptosomes were lysed with the lysis buffer supplemented with protease inhibitor cocktail (Roche). The co-IP was performed by incubating the 3E9 antibody with the synaptosomal lysate overnight at 4°C, followed by 4 hr incubation with protein G beads. Protein bound beads were stringently washed, boiled in 2x sample loading buffer and the supernatants were used for LC-MS/MS analysis. The proteins were separated on a 12% SDS-PAGE and subjected to in-gel digestion using trypsin. The tryptic peptides were separated and analyzed using a Dionex Ultimate 3000 RSLCnano system coupled to a Q Exactive instrument (Thermo Fisher Scientific, San Jose, CA) as previously described (Gallart-Palau et al., 2019). Raw data files were converted to mascot generic file format using Proteome Discoverer 1.4 (Thermo Fisher Scientific, San Jose, CA). The MS/MS spectra were subjected to Mascot database search against UniProt mouse protein database. The exponentially modified protein abundance index (emPAI) of identified individual proteins were used for label-free protein quantitation as previous described (Gallart-Palau et al., 2019). Imp-β1 specific binding proteins determined by subtracting the Imp-β1 co-IP proteins with the background proteins in the IgG control sample were shortlisted for further analysis.

