## Supplementary Materials for "LOCAL REGULATION AND FUNCTION OF IMPORTIN-β1 IN HIPPOCAMPAL NEURONS DURING TRANSCRIPTION-DEPENDENT PLASTICITY"

**Supplementary Figure 1.**

**(A)** Lysate from transfected HEK 293T cells with HA-tagged human Imp-β1 plasmids and lysates from Imp-β1 IP experiments in non-transfected HEK 293T cells were probed with antibodies targeted against Imp-β1, HA protein tag and GAPDH. **(B)** Representative confocal micrographs of dentate gyrus neurons in basal and depolarized (cLTP) acute forebrain slices. Depolarized acute forebrain slices were treated with forskolin (40 µM, 15 min) and KCl (50 mM, 10 min) in $Mg^{2+}$ free aCSF before slices were fixed and immunohistochemistry performed with antibodies targeted against MAP2 (cyan), CRTC1 (red), Imp-β1 (green) and Hoechst nuclear dye (blue). The nuclear-to-cytoplasmic ratio of CRTC1 and Imp-β1 were quantified and the group data from independent experiments were plotted. Unpaired T-test analyses were conducted between stimulated and basal condition (N=3; ***$p$ < 0.001, ****$p$ < 0.0001). Scale bar, 20 µm.

**Supplementary Figure 2.**

**(A)** Hippocampal neurons were treated with glutamate (Glut; 40 µM, 5 min) with or without INI43 (5 µM, 1 h) before fixing and immunolabeling with antibodies targeted against MAP2 (cyan), Imp-β1 (red) and Hoechst nuclear dye (blue in merged image). The Imp-β1 nuclear-to-cytoplasmic ratio was quantified and the group data from independent experiments were plotted. Statistical analyses performed with one way-ANOVA with Dunn's post-hoc analyses (N=2; *$p$ < 0.05, ****$p$ < 0.0001). Scale bar, 10 µm. **(B)** Representative confocal micrographs of TTX-silenced and KCl-depolarized cultured hippocampal neurons (DIV14) transfected with sGFP-NFATc3 plasmids on DIV13, with or without INI43 treatment, immunolabeled with antibodies targeted against MAP2 (cyan), sGFP-NFATc3 (red) and Hoechst nuclear dye (blue in merged image). All neurons were pre-treated with TTX (1 µM, 30 min) in Tyrode's solution to reduce baseline activity. INI43 (5 µM, 30 min) treatment was performed where indicated together with TTX pretreatment. Neurons were then depolarized with KCl (50 mM, 3 min) in the presence or absence of INI43. All the neurons were recovered in Tyrode's solution containing TTX (1 µM, 10 min), with INI43 (5 µM) where indicated. The nuclear-to-cytoplasmic ratio of sGFP-NFATc3 staining in neurons were quantified and group data is plotted. Statistical analyses performed on group data use one way-ANOVA with Dunn's post-hoc analyses (N=2; *$p < 0.05$, **$p < 0.05$, ****$p < 0.0001$). Scale bar, 5 µm. **(C)** Representative confocal micrographs of basal and stimulated cultured hippocampal neurons, in the presence or absence of INI43, immunolabeled with antibodies targeted against MAP2 (cyan), CRTC1 (red) and Hoechst nuclear dye (blue in merged image). Stimulated cultured neurons were treated with bicuculline (Bic; 50 µM, 30 min) + 4AP (200 µM, 30 min), with or without INI43 (5 µM, 1 h). For experiments with INI43, neurons were pre-treated with the inhibitor prior to stimulation. The nuclear-to-cytoplasmic ratio of CRTC1 in neurons were quantified and group data is plotted. Statistical analyses performed on group data use one way-ANOVA with Dunn's post-hoc analyses (N=2; ****$p < 0.0001$). Scale bar, 10 µm.  **(D)** Mean Ct values of the housekeeping gene, HPRT1, is depicted in bar graph for neuronal lysates obtained for quantitative PCR (qPCR) analyses. Cultured neurons were treated with various pharmacological agents including bicuculline (Bic; 50 µM, 30 min) + 4AP (200 µM, 30 min) and INI43 (5 µM, 1 h). For experiments with INI43, neurons were pre-treated with the inhibitor prior to and during stimulation. **(D-ii)** Representative confocal micrographs of basal neurons, with or without INI43 treatment, as well as $H_2O_2$ treated neurons immunolabeled with MAP2 (cyan), cleaved caspase 3 (red) and Hoechst nuclear dye (blue in merged image). Cultured neurons were treated with various pharmacological agents including INI43 (5 µM, 1 h) and $H_2O_2$ (500µM, 2 h). The mean cleaved caspase 3 staining in the soma were quantified and normalized against the neurons without INI43 treatment. The group data is plotted. Statistical analyses performed with one way-ANOVA with Dunn's post-hoc analyses (N=2; ***$p < 0.001$, ****$p < 0.0001$). Scale bar, 2 µm.

**Supplementary Figure 3.**

Rat cultured hippocampal neurons expressing Dendra2 constructs were photoconverted using UV (405 nm) laser and tracked for 15 min (1 frame / min). Time lapse images of the somatic photoconverted Dendra2 signals (every $5^{th}$ frame) are shown for basal and stimulated neurons. Neurons were imaged either at basal state or bicuculline-treated (40 μm) in $Mg^{2+}$ free Tyrode's solution.

**Supplementary Figure 4.**

**(A-i)** Illustration for puromycin labeled proximity ligation assay (Puro-PLA). Cycloheximide (CHX; 355 µM, 30 min) is first added to the cultured hippocampal neurons to stall translation at the site of synthesis. Puromycin (Puro; 3 µM, 15 min) is then added to label the truncated nascent peptides. When the conjugated antibodies targeted against puromycin ($PLA^{minus}$ ; red Y) and Imp-β1 (3E9) ($PLA^{plus}$ ; blue Y) are in close proximity, the template linker oligo will ligate both the plus and minus probes into a circle, enabling circling amplification. Lastly, the binding of the fluorescent molecules on the amplified probes generates the PLA signal (green fluorescence). **(A-ii)** Cultured mouse hippocampal neurons were pre-treated with cycloheximide (CHX; 50 µM, 30min) before Puro-PLA, in the presence of single or double probes before being fixed and immunolabeled with antibodies targeted against MAP2 (violet) and Hoechst nuclear dye (blue in merged image). The puro-PLA signal specific for Imp-β1 appears as green puncta. The integrated Puro-PLA intensity normalised to total MAP2 area in the field was quantified and the group data from independent experiments were plotted. Statistical analyses performed on group data use one way-ANOVA with Dunn's post-hoc analyses (N=2, n=20; $*p < 0.05$, $**p < 0.01$). Scale bar, 2 µm.

**Supplementary Figure 5.**

**(A)** Cultured mouse hippocampal neurons stimulated with KCl (50 mM, 5 min) and glycine (100 µM, 5 min), tetrodotoxin (TTX; 1 µM, 1 h) and APV (40 µM, 1 h), before being fixed and immunolabeled with antibodies targeted against MAP2 (violet) and Imp-β1 (red). The normalized dendritic Imp-β1 staining against MAP2 staining were quantified and the group data is plotted and indicated in the table. Statistical analyses performed on group data use one way-ANOVA with Dunn's post-hoc analyses (*$p$ <0.05, ****$p$ <0.0001). **(B)** Cultured hippocampal neurons were treated with and without N-ethylmaleimide (NEM; 50 µM, 10 min) in Tyrode's solution and lysosomes from both conditions were labeled with Lysotracker (1 µM, 10 min). The movement of the lysosomes were then imaged using live confocal microscopy (1s / frame for 60 s) and tracked using particle displacement module (IMARIS). The lines were color coded based on individual lysosomal displacement length. The cumulative displacement of the five fastest moving lysosomes for each experiment (N=3; n=15) were plotted against time along with the average displacement of all the lysosomes captured in each condition. Unpaired T-test analysis was conducted between NEM-treated and mock condition (N=3, mock: n=2971, NEM: n=2393; ****$p$ < 0.0001). **(C)** The mean MAP2 intensity of dendrites from neurons treated with BDNF- (50 ng/mL, 20 min), DHPG- (100 µM, 5 min with 15 min recovery) and/or cycloheximide (CHX; 50 µM, 30 min) were plotted. All neurons were pre-treated with NEM (50 µM, 10 min) and MG132 (5 µM, 30 min) before stimulation. Statistical analyses performed on group data use one way-ANOVA with Dunn's post-hoc analyses (N=2).

**Supplementary Figure 6.**

(A) Different fractions from the synaptosomal extraction protocol were harvested, subjected to SDS-PAGE and probed with antibodies targeted against PSD95, synaptophysin (SYPH), Histone-H3, and Imp-β1. Lys, forebrain homogenate; P1: nuclear fraction; P2 and P3 pellets; S1 and S2: supernatants harvested during enrichment; Syn, purified synaptosomes fraction. The fold change of the synaptic markers in the purified synaptosomes were quantified and normalized against the forebrain homogenate (Lys) and the group data represented as bar graph (SYPH: N=3; PSD95: N=5). **(B)** Synaptosomes in basal and stimulated (glycine, 100 µM, 5 min followed by KCl, 50 mM, 5 min) states were probed with antibodies targeted against CRTC1. The ratio of dephosphorylated to total CRTC1 were quantified and plotted on a before-after graph (N=3). **(C-i)** A schematic flowchart on the isolation of candidate proteins based on subcellular localization using various synaptic and nuclear databases. At each step, the number of proteins in the candidate list is stated in brackets. The number of proteins found at least once in the three independent synaptic databases (199/277), MGI nuclear database (105/277) or belonging to both group (83/277) are listed. **(C-ii)** GO analyses using DAVID showing the top enriched GO terms for molecular functions (MF) and biological processes (BP) from the candidate list (277 protein). **(D)** Lysates from transfected HEK 293T cells with Flag-tagged Imp-α1-3 and 6, as well as paGFP-tagged Imp-α4 plasmids were probed with antibodies targeted against individual Imp-α isoform. Red arrows indicate the overexpressed Imp-α. **(E)** Cultured mouse hippocampal neurons treated with or without leptomycin B (LMB; 20 nM, 3 h) before fixing and immunolabeling with antibodies targeted against MAP2 (cyan), NDRG1 (green) and Hoechst nuclear dye (blue in merged image). The nuclear-to-cytoplasmic ratio of NDRG1 were quantified and the group data from independent experiments were plotted. Unpaired T-test analyses were conducted between mock and LMB treated groups (N=2; ****$p < 0.0001$). Scale bar, 2 µm.

**Supplementary Figure 7.**

83 candidate proteins shortlisted that has an enhanced association with Imp-β1 following synaptic stimulation in purified synaptosomes. The emPAI scores are reflected in each condition. Y – Evidence on nuclear shuttling in response to stimulus in non-neuronal cells. **Y** – Evidence on nuclear shuttling in response to stimulus in neurons. **Y** – Known synapse-to-nucleus signaling proteins.

Supplementary Fig. 1

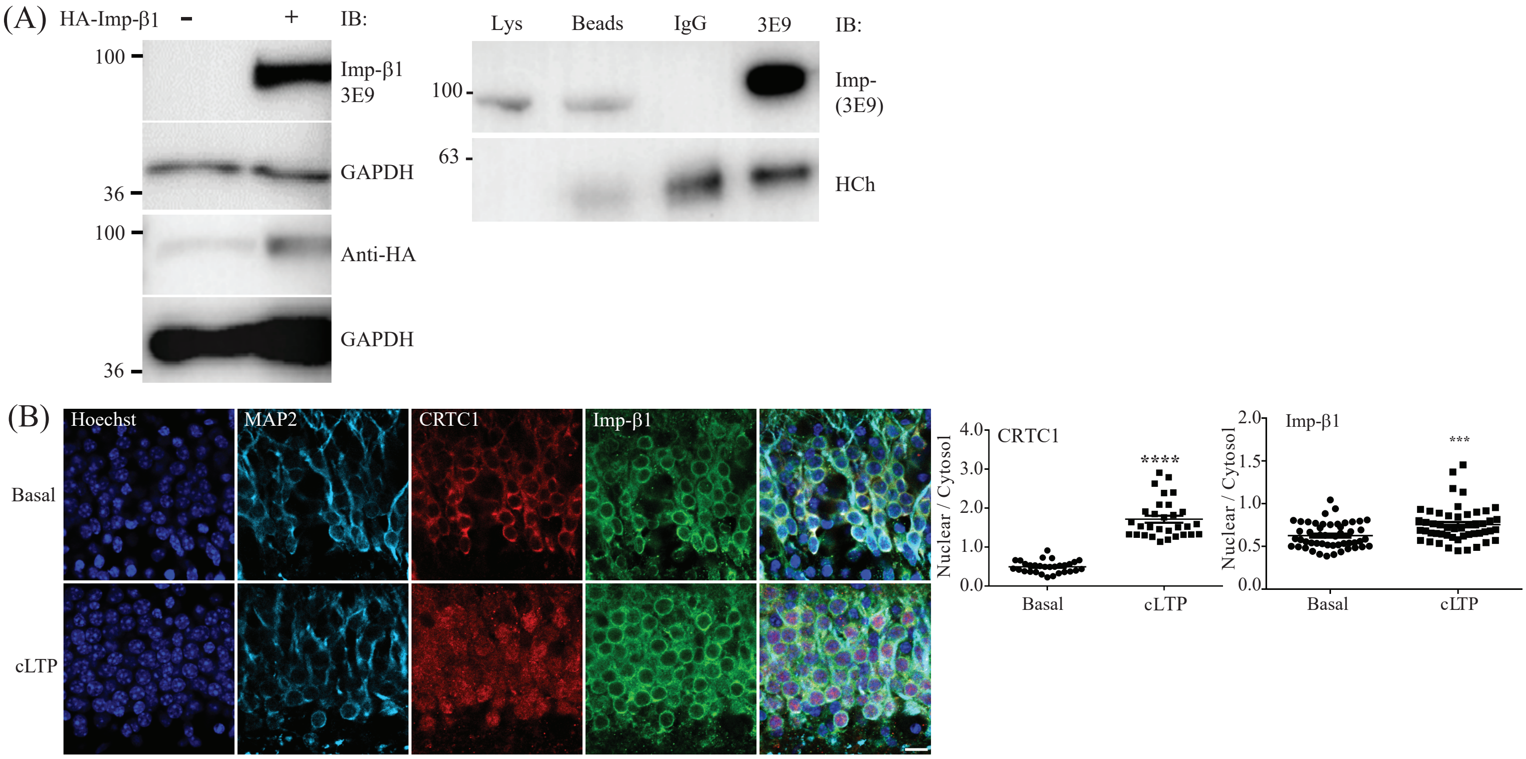

Supplementary Fig. 2

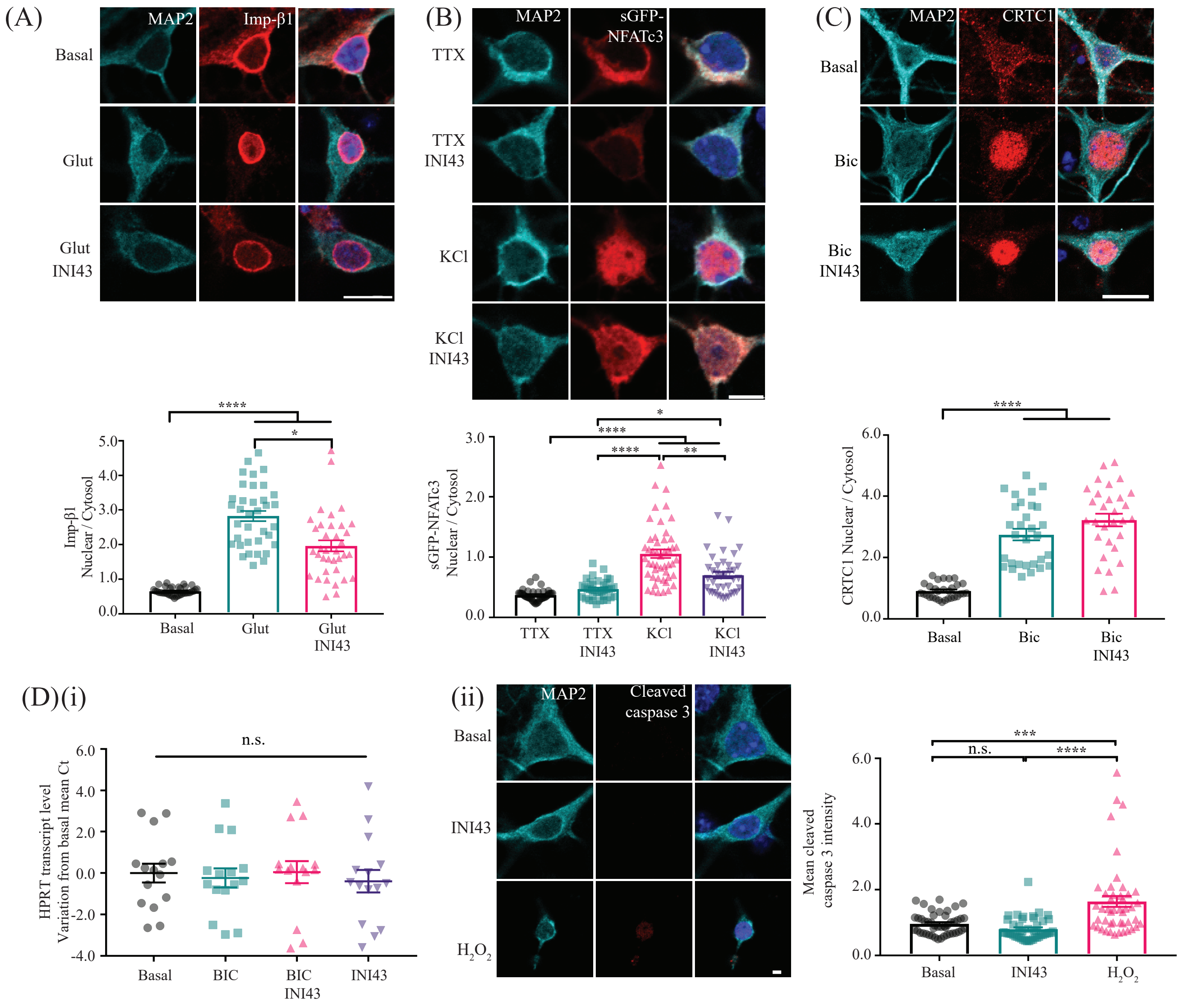

### Supplementary Figure 3.

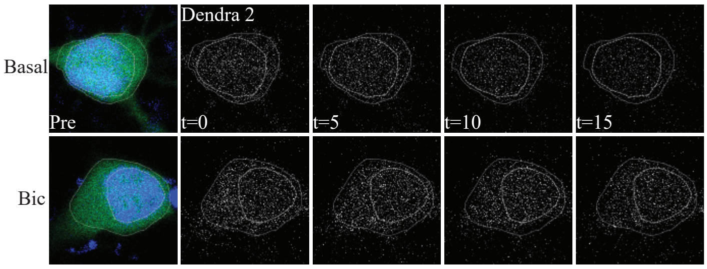

### Supplementary Figure 4.

(A)(i)

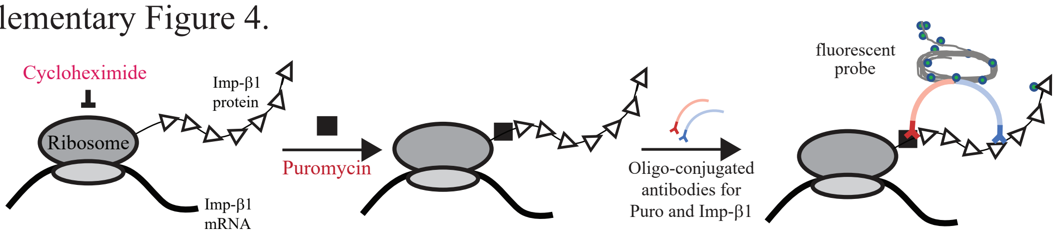

(ii)

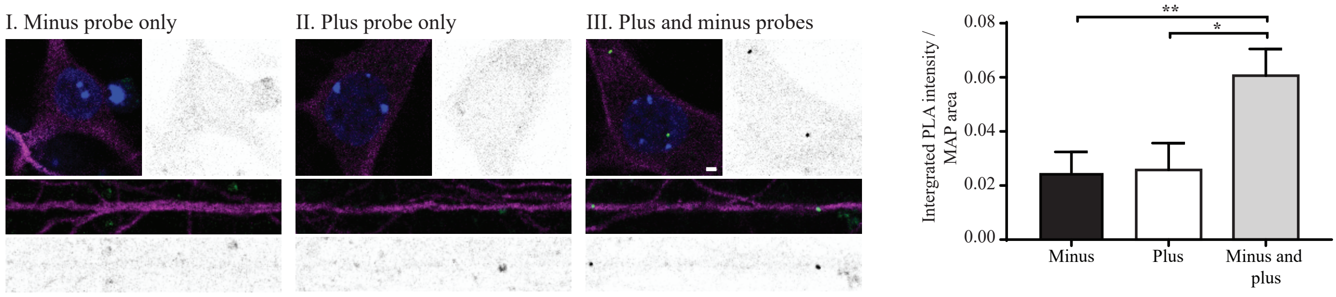

### Supplementary Figure 5.

(A)

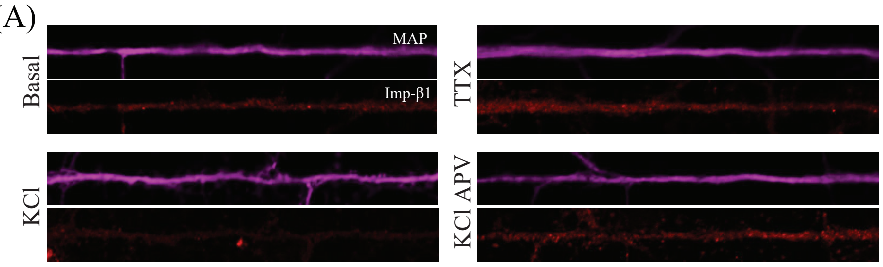

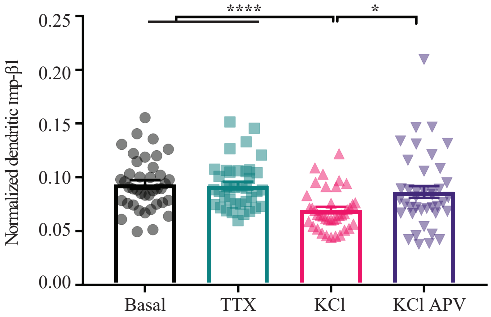

| | Basal | TTX | KCl | KCl APV |
|---|---|---|---|---|
| **Sample size (N=3)** | 40 | 38 | 39 | 40 |
| **Imp-β1 / MAP2 ± SEM** | 0.0935 ± 0.00385 | 0.0925 ± 0.00356 | 0.0697 ± 0.00301 | 0.0864 ± 0.00553 |

(B)

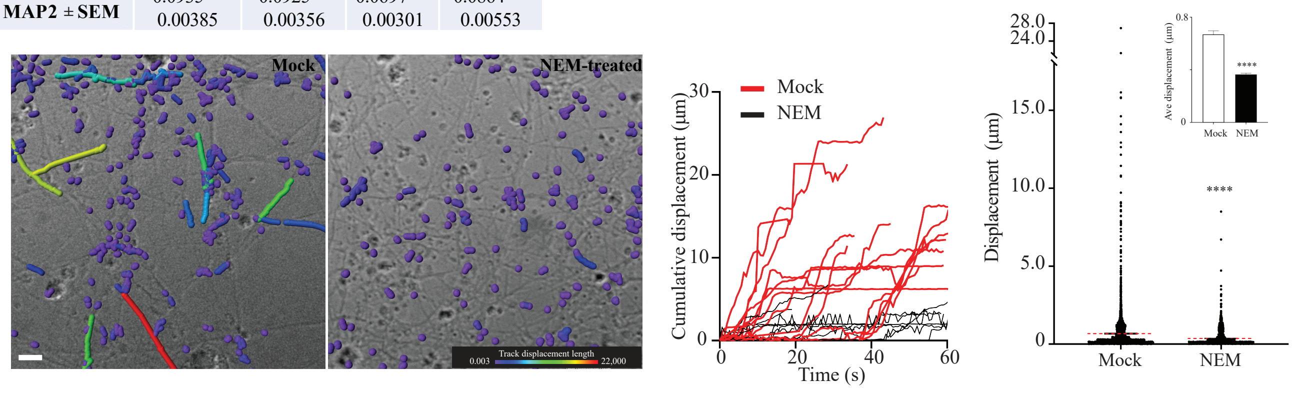

(C)

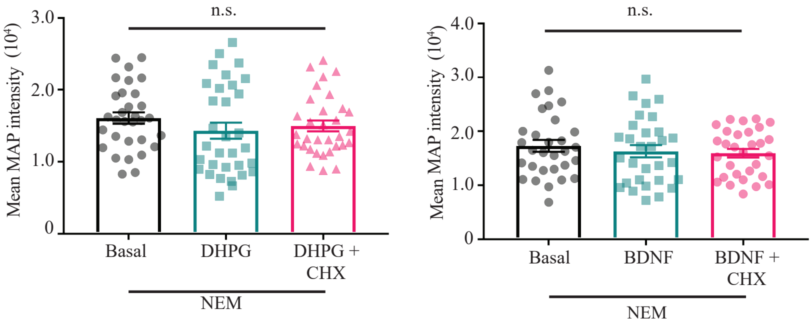

Supplementary Figure 6.

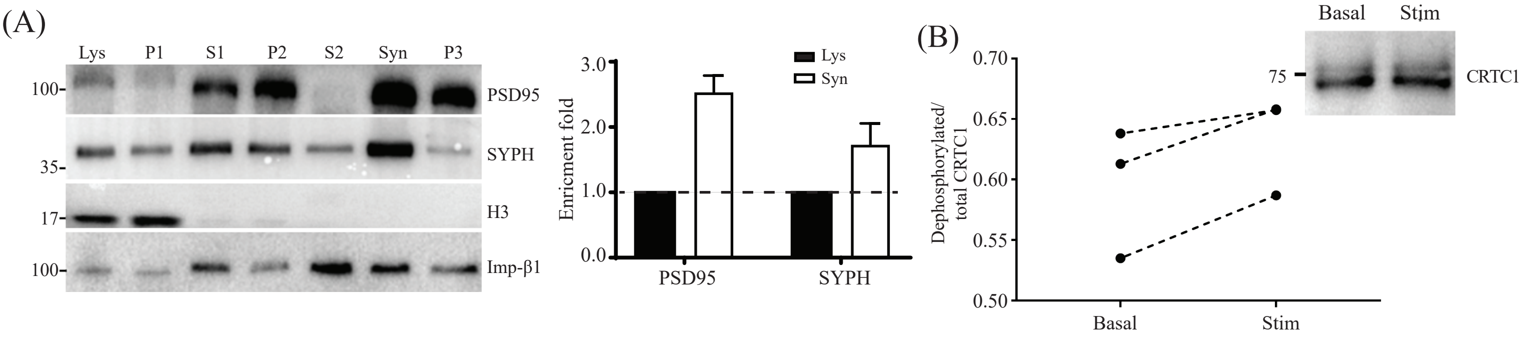

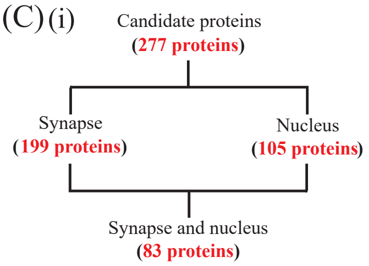

(ii)

| Annotation Cluster 1 | Enrichment: 11.05 | p-values |
|---|---|---|
| GOTERM_ CC | cell-cell adherens junction | 5.1 x $10^{-14}$ |
| GOTERM_ MF | cadherin-binding involved in cell-cell adhesion | 2.3 x $10^{-13}$ |
| GOTERM_ BP | cell-cell adhesion | 6.4 x $10^{-8}$ |

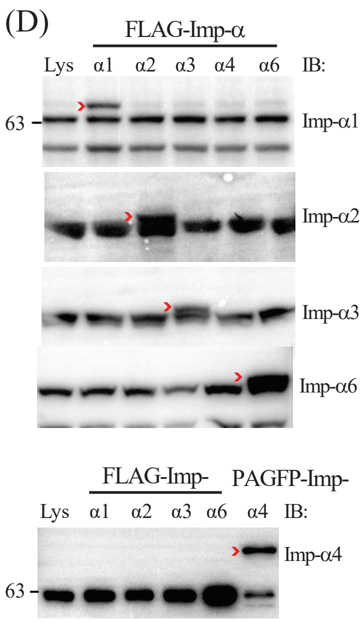

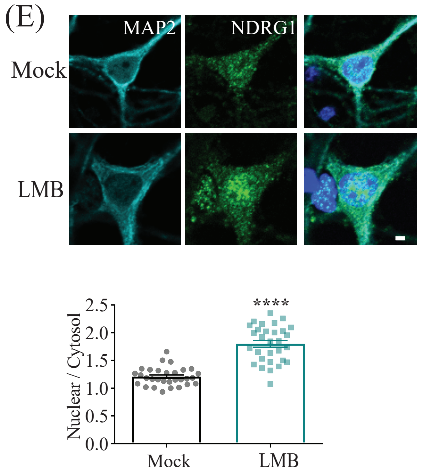

)

**Supplementary Figure 7.**

| | | | Stimulated | | | Basal | | | | | |
|---|---|---|---|---|---|---|---|---|---|---|---|
| **Accession** | **GS** | **Name** | **IgG** | **Imp-β1** | **B1/IgG** | **IgG** | **Imp-β1** | **B1/IgG** | **Stimulated / Basal** | **Nuclear shuttling** | **Remarks** |
| **Q3TED3** | Acly | ATP-citrate synthase | -1 | 0.44 | -0.44 | -1 | 0.21 | -0.21 | 2.1 | | |
| **D3Z0T1** | Add1 | Alpha-adducin | -1 | 0.28 | -0.28 | -1 | -1 | 1 | -0.28 | Y | Contains NLS, calcium-dependent nuclear shuttling in cancer cells (Kiang & Leung, 2018) |
| **A0A1D5RMG4** | Agap3 | Arf-GAP with GTPase, ANK repeat and PH domain-containing protein 3 | -1 | 0.44 | -0.44 | -1 | 0.04 | -0.04 | 11 | Y | Its splice variant, CRAG, contains nuclear localization signal. ROS triggered the association of CRAG with polyQ and the nuclear translocation of CRAG-polyQ complex (Qin et al., 2006). CRAG knock out mice exhibited suppressed kainic acid induced cfos expression in hippocampus. (Nagashima et al., 2019) |
| **A2ALT5** | Ahcy | Adenosylhomocysteinase | -1 | 0.44 | -0.44 | -1 | -1 | 1 | -0.44 | Y | Nucleocytoplasmic shuttling in basal Hela cells (Grbeša et al., 2017) Nuclear accumulation in transcriptionally active cells (Radomski, Kaufmann, & Dreyer, 1999) |
| **Q9CZS1** | Aldh1b1 | Aldehyde dehydrogenase X, mitochondrial | -1 | 0.11 | -0.11 | -1 | -1 | 1 | -0.11 | | |
| **S4R2J3** | Anks1b | Ankyrin repeat and sterile alpha motif domain-containing protein 1B (Fragment) | -1 | 0.23 | -0.23 | -1 | -1 | 1 | -0.23 | **Y** | Contains monopartite NLS, Activity-dependent synapse-to-nucleus shuttling. (Jordan, |

| | | | | | | | | | | | |
|---|---|---|---|---|---|---|---|---|---|---|---|
| | | (AIDA) | | | | | | | | | Fernholz, Khatri, & Ziff, 2007) |
| **Q3TWT5** | Asah1 | Acid ceramidase | -1 | 0.44 | -0.44 | -1 | -1 | 1 | -0.44 | | |
| **D3Z2Y8** | Bdh1 | D-beta-hydroxybutyrate dehydrogenase, mitochondrial | -1 | 0.15 | -0.15 | -1 | 0.12 | -0.12 | 1.25 | | |
| **Q6P1B9** | Bin1 | Bin1 protein | -1 | 0.5 | -0.5 | -1 | -1 | 1 | -0.5 | | |
| **A0A0M3HEP6** | Cadps2 | Calcium-dependent secretion activator 2 | -1 | 0.12 | -0.12 | -1 | 0.06 | -0.06 | 2 | | |
| **F8WHB5** | Camk2a | Calcium/calmodulin-dependent protein kinase type II subunit alpha | -1 | 0.16 | -0.16 | -1 | -1 | 1 | -0.16 | Y | Contains NLS, NMDAR dependent PSD localization (nuclear exit) (Zalcman, Federman, & Romano, 2018) |
| **O70589-2** | Cask | Isoform 2 of Peripheral plasma membrane protein CASK | -1 | 0.44 | -0.44 | -1 | -1 | 1 | -0.44 | Y | Calcium dependent nuclear shuttling (Ojeh, Pekovic, Jahoda, & Määttä, 2008; Schuh, Uldrijan, Gambaryan, Roethlein, & Neyses, 2003) |
| **G3UXG7** | Csnk2b | Casein kinase II subunit beta | -1 | 0.16 | -0.16 | -1 | -1 | 1 | -0.16 | Y | Contains NLS. FGF2 binding targeted the holoenzyme containing Csnk2b into the nucleus (Filhol et al., 2003). Mitogenic stimulation induces nuclear translocation (Lorenz, Pepperkok, Ansorge, & Pyerin, 1993). |
| **A0A0G2JDL3** | Cdk5 | Cyclin-dependent-like kinase 5 | -1 | 0.13 | -0.13 | -1 | -1 | 1 | -0.13 | **Y** | Nuclear translocation in depolarized neurons (Z. Liang et al., 2015) Stress induces nuclear translocation in SHSY5Y cells (Song et al., 2016) |
| **F6QFL0** | Chchd3 | MICOS complex subunit | -1 | 0.32 | -0.32 | -1 | -1 | 1 | -0.32 | | |
| **Q3UPG6** | Clint1 | Clathrin interactor 1 | -1 | 0.13 | -0.13 | -1 | -1 | 1 | -0.13 | | |
| **Q8C5W0-4** | Clmn | Isoform 4 of Calmin | -1 | 0.44 | -0.44 | -1 | -1 | 1 | -0.44 | | |

| | | | | | | | | | | |
|---|---|---|---|---|---|---|---|---|---|---|
| **Q3TEA5** | Cops3 | COP9 signalosome complex subunit 3 | -1 | 0.44 | -0.44 | -1 | -1 | 1 | -0.44 | | |
| **Q3UZT7** | Ctnnb1 | Catenin beta-1 | -1 | 0.28 | -0.28 | -1 | -1 | 1 | -0.28 | **Y** | NMDAR dependent synapse-to-nucleus translocation (Schmeisser, Grabrucker, Bockmann, & Boeckers, 2009)<br>NMDAR dependent activation of calpain induced cleavage of CTNNB1 at N-terminus, which in turns, accumulates in the nucleus in hippocampal neurons (Abe & Takeichi, 2007) |
| **B7ZNF6** | Ctnnd2 | Catenin delta-2 | -1 | 0.21 | -0.21 | -1 | -1 | 1 | 0.21 | | Contains NLS (Koutras & Lévesque, 2011)<br>Nuclear translocation required for NPRAP gene regulation (Koutras, Lessard, & Lévesque, 2011) |
| **Q9D0M3-2** | Cyc1 | Isoform 2 of Cytochrome c1, heme protein, mitochondrial | -1 | 0.15 | -0.15 | -1 | 0.05 | -0.05 | 3 | | |
| **Q56A15** | Cycs | Cytochrome c | -1 | 0.44 | -0.44 | -1 | -1 | 1 | -0.44 | Y | Apoptotic signal, DNA damage, depletion of VDAC1 during differentiation and nitro-oxidative stress (for a protective role) drives nuclear accumulation of Cycs (González-Arzola et al., 2019) |
| **D3YZY5** | Dhodh | Dihydroorotate dehydrogenase (quinone), mitochondrial | -1 | 0.44 | -0.44 | -1 | -1 | 1 | -0.44 | | |
| **B1AXY1** | Dnaja1 | DnaJ homolog subfamily A member 1 | -1 | 0.16 | -0.16 | -1 | -1 | 1 | -0.16 | Y | Mutant glucocorticoid receptor results in nuclear accumulation of Dnaja1 in |

| | | | | | | | | | | | |
|---|---|---|---|---|---|---|---|---|---|---|---|
| | | | | | | | | | | | Cos cells (Tang, Ramakrishnan, Thomas, & DeFranco, 1997)<br>Dj2 accumulates in nucleolus with reduced staining in cytosol following prolong heat treatment (Davis, Alevy, Chellaiah, Quinn, & Mohanakumar, 1998)<br>Dj1 accumulates in nucleoli after heat shock (Terada & Mori, 2000) |
| **D3Z6U8** | FMR1 | Fragile X mental retardation protein 1 homolog | -1 | 0.15 | -0.15 | -1 | 0.08 | -0.08 | 1.66 | | Contains NLS and NES, known nucleocytoplasmic shuttling protein that shuttles mRNA out of the nucleus (Feng et al., 1997). |
| **Q00612** | G6pdx | Glucose-6-phosphate 1-dehydrogenase X | -1 | 0.11 | -0.11 | -1 | -1 | 1 | -0.11 | | |
| **A0A0G2JGS8** | Gabrb1 | Gamma-aminobutyric acid receptor subunit beta-1 (Fragment) | -1 | 0.13 | -0.13 | -1 | -1 | 1 | -0.13 | | |
| **Z4YKV1** | Gnas | Guanine nucleotide-binding protein G(s) subunit alpha isoforms short | -1 | 0.28 | -0.28 | -1 | -1 | 1 | -0.28 | | |
| **F7ALS6** | Got1 | Aspartate aminotransferase, cytoplasmic | -1 | 0.23 | -0.23 | -1 | -1 | 1 | -0.23 | | |
| **A0A0U1RQ18** | Gpi1 | Glucose-6-phosphate isomerase | -1 | 0.28 | -0.28 | -1 | -1 | 1 | -0.28 | Y | Hypoxia drives nuclear import of GPI in cancer cells (Funasaka, Yanagawa, Hogan, & Raz, 2005) |
| **Q60631** | Grb2 | Growth factor receptor-bound protein 2 | -1 | 0.14 | -0.14 | -1 | -1 | 1 | -0.14 | Y | Accumulates in the nucleus and plasma membrane after EGF stimulation with a reduction in cytosol in |

| | | | | | | | | | | | |
|---|---|---|---|---|---|---|---|---|---|---|---|
| | | | | | | | | | | | A431 cells (Yamazaki et al., 2002) |
| **Q5KU03** | Gsk3b | Glycogen synthase kinase 3 beta | -1 | 0.28 | -0.28 | -1 | -1 | 1 | -0.28 | Y | Cell senescence (Zmijewski & Jope, 2004), pro-apoptotic stimuli (Bijur & Jope, 2001) induce nuclear translocation. Contains bipartite NLS, growth factor stimulation induces nuclear export while cell cycle progression induces nuclear import (Meares & Jope, 2007) |
| **Q8BG05** | Hnrnpa3 | Heterogeneous nuclear ribonucleoprotein A3 | -1 | 0.11 | -0.11 | -1 | -1 | 1 | -0.11 | | Nucleocytoplasmic shuttling protein for RNA export (Papadopoulou, Boukakis, Ganou, Patrinou-Georgoula, & Guialis, 2012) |
| **P61979** | Hnrnpk | Heterogeneous nuclear ribonucleoprotein K | -1 | 0.22 | -0.22 | -1 | -1 | 1 | -0.22 | Y | Hnrnpk relocalized to cytoplasm during VSV infection (Pettit Kneller, Connor, & Lyles, 2009). Serum stimulation or activated pERK drives phosphorylated Hnrnpk into cytosol (Habelhah et al., 2001). BDNF activity drives HnrnpK dendritic accumulation in hippocampal neurons (Morais, 2014). |
| **A0PJ91** | Hsp90aa1 | Hsp90aa1 protein | -1 | 0.44 | -0.44 | -1 | -1 | 1 | -0.44 | Y | HSP90 interacts with Imp-β1 for glucocorticoid receptor nuclear import (Echeverría et al., 2009) B-napthoflavone induces aryl hydrocarbon receptor |
| **E9Q3D6** | Hsp90ab 1 | Heat shock protein HSP 90-beta | -1 | 0.44 | -0.44 | -1 | -1 | 1 | -0.44 | | |

| | | | | | | | | | | |
|---|---|---|---|---|---|---|---|---|---|---|
| | | | | | | | | | | complexing with HSP90 to enter nucleus in HeLa cells (Tsuji et al., 2014) Hypoxia drives nuclear entry of HSP90 (Katschinski et al., 2004) |
| **A0A0J9YTZ7** | Hsph1 | Heat shock protein 105 kDa (Fragment) | -1 | 0.32 | -0.32 | -1 | -1 | 1 | -0.32 | Y | The β-splice variant of Hsph1 shuttles to the nucleus following stress to induce HSP70 expression (Stetler et al., 2010). Nuclear entry dependent on Imp-β1 cargo (Saito, Yamagishi, & Hatayama, 2007). |
| **A2AFQ0** | Huwe1 | E3 ubiquitin-protein ligase HUWE1 | -1 | 0.44 | -0.44 | -1 | -1 | 1 | -0.44 | | |
| **Q9EPL8** | Ipo7 | Importin-7 | -1 | 0.42 | -0.42 | -1 | 0.05 | -0.05 | 8.4 | **Y** | |
| **F6U8U1** | Itsn1 | Intersectin-1 | -1 | 0.44 | -0.44 | -1 | -1 | 1 | -0.44 | | LMB traps Itsn1 in the nucleus, Imp-α cargo (Alvisi et al., 2018) |
| **Q02257** | Jup | Junction plakoglobin | -1 | 0.28 | -0.28 | -1 | -1 | 1 | -0.28 | Y | Inadequate MAPK activity results in APC cytoplasmic sequestration along with Jup nuclear accumulation in cancer cells (Hildesheim, Salvador, Hollander, & Fornace, 2005) |
| **Q60960** | Kpna1 | Importin subunit alpha-5 | -1 | 0.23 | -0.23 | -1 | 0.05<br>5 | -0.055 | 4.09 | Y | |
| **O35343** | Kpna4 | Importin subunit alpha-3 | -1 | 0.29 | -0.29 | -1 | 0.06 | 0.13 | -2.26 | Y | |
| **Q4FJZ2** | Kpna6 | Importin subunit alpha-6 | -1 | 0.27 | -0.27 | -1 | 0.15 | -0.15 | 1.74 | Y | |
| **Q3TFE8** | Kpnb1 | Importin subunit beta-1 | -1 | 1.16 | -1.16 | -1 | 0.93 | -0.93 | 1.25 | Y | |
| **D3Z3G6** | MAPK3 | Mitogen-activated protein kinase | -1 | 0.15 | -0.15 | -1 | 0.02 | -0.02 | 7.5 | **Y** | Synapse-to-nucleus signaling proteins dependent on synaptic activation (Zhai, Ark, Parra-Bueno, & Yasuda, 2013). |

| | | | | | | | | | | | |
|---|---|---|---|---|---|---|---|---|---|---|---|
| | | | | | | | | | | | Nuclear entry dependent on Imp-β1 (Perlson et al., 2006) and IPO7 (Lorenzen et al., 2001). |
| **Q3UH19** | Mapt | Microtubule-associated protein | -1 | 0.68 | -0.68 | -1 | -1 | 1 | -0.68 | Y | Contains NLS (Alonso et al., 2010)<br>Disrupted microtubules (Siano et al., 2019), oxidative and heat stress (Sultan et al., 2011) induce nuclear translocation of Tau in neurons. |
| **E9PVF3** | Ndrg1 | N-myc downregulated gene 1 | -1 | 0.21 | -0.21 | -1 | -1 | 1 | -0.21 | Y | DNA damage drives nuclear translocation of NDRG1 in p53 dependent manner (Kitowska & Pawełczyk, 2010)<br>Hypoxia induces nuclear accumulation of NDRG1 in human trophoblasts (Shi, Larkin, Chen, & Sadovsky, 2013) |
| **Q80TQ3** | Nefh | Neurofilament heavy polypeptide | -1 | 0.44 | -0.44 | -1 | -1 | 1 | -0.44 | | |
| **F6V7K3** | Npepps | Puromycin-sensitive aminopeptidase | -1 | 0.12 | -0.12 | -1 | -1 | 1 | -0.12 | | |
| **A2AT02** | Nsfl1c | NSFL1 cofactor p47 | -1 | 0.44 | -0.44 | -1 | -1 | 1 | -0.44 | | |
| **E0CXD4** | Pcdh1 | Protocadherin-1 | -1 | 0.28 | -0.28 | -1 | 0.08 | -0.08 | 3.5 | Y | ICD of α-Pcdh and γ-Pcdh undergo nuclear shuttling in cell lines (Haas, Frank, Véron, & Kemler, 2005; Hambsch, Grinevich, Seeburg, & Schwarz, 2005)<br>Processing of human Pcdh (Fat1) releases C-terminal that shuttles into the nucleus in cell lines (Magg, Schreiner, Solis, Bade, & Hofer, 2005) |

| | | | | | | | | | | |
|---|---|---|---|---|---|---|---|---|---|---|
| **F2Z4A5** | Pdpk1 | 3-phosphoinositide-dependent protein kinase 1 | -1 | 0.26 | -0.26 | -1 | -1 | 1 | -0.26 | Y | Contains NLS. IGF1, insulin, nerve growth factor and activated src kinase stimulates nuclear localization of Pdpk1. PI3K signaling modulates nucleocytoplasmic regulation of Pdpk1. (Kikani, Dong, & Liu, 2005; Lim, Kikani, Wick, & Dong, 2003; Scheid, Parsons, & Woodgett, 2005) Stimulated EP4 receptors by prostaglandins drives nuclear translocation of Pdpk1 in human eosinophils (Sturm et al., 2015) |
| **A2AS44** | Pkp4 | Plakophilin-4 | -1 | 0.28 | -0.28 | -1 | -1 | 1 | -0.28 | | |
| **Q9Z2M7** | Pmm2 | Phosphomannomutase 2 | -1 | 0.12 | -0.12 | -1 | -1 | 1 | -0.12 | | |
| **A0A087WRA7** | Ppp1r7 | Protein phosphatase 1 regulatory subunit 7 | -1 | 0.25 | -0.25 | -1 | -1 | 1 | -0.25 | | |
| **Q9QUM9** | Psma6 | Proteasome subunit alpha type-6 | -1 | 0.13 | -0.13 | -1 | -1 | 1 | -0.13 | | |
| **Q9Z2U0** | Psma7 | Proteasome subunit alpha type-7 | -1 | 0.12 | -0.12 | -1 | -1 | 1 | -0.12 | | Interacts with Imp-α2 (Arjomand et al., 2014) |
| **Q9CWH6** | Psma8 | Proteasome subunit alpha type-7-like | -1 | 0.18 | -0.18 | -1 | -1 | 1 | -0.18 | | |
| **F2Z3U3** | Raph1 | Ras association (RalGDS/AF-6) and pleckstrin homology domains 1 | -1 | 0.44 | -0.44 | -1 | -1 | 1 | -0.44 | | |
| **A0A1B0GSG5** | Rnh1 | Ribonuclease inhibitor | -1 | 0.44 | -0.44 | -1 | -1 | 1 | -0.44 | Y | Cellular stress induces nuclear accumulation of Rnh1 in HeLa cells (Pizzo et al., 2013) |
| **Q3TDK6** | Rogdi | Protein rogdi homolog | -1 | 0.13 | -0.13 | -1 | -1 | 1 | -0.13 | | Suggested as presynaptic to nucleus shuttling protein |

| | | | | | | | | | | | |
|---|---|---|---|---|---|---|---|---|---|---|---|
| | | | | | | | | | | | (Riemann, Wallrafen, & Dresbach, 2017) |
| **P62830** | Rpl23 | 60S ribosomal protein L23 | -1 | 0.41 | -0.41 | -1 | -1 | 1 | -0.41 | | |
| **D3YVM5** | Rplp0 | 60S acidic ribosomal protein P0 (Fragment) | -1 | 0.23 | -0.29 | -1 | -1 | 1 | -0.29 | | |
| **P62301** | Rps13 | 40S ribosomal protein S13 | -1 | 0.49 | -0.49 | -1 | -1 | 1 | -0.49 | | |
| **S4R223** | Rps19 | 40S ribosomal protein S19 | -1 | 0.54 | -0.54 | -1 | 0.22 | -0.22 | 2.49 | | |
| **A0A0N4SWD0** | Scrn1 | Secernin-1 (Fragment) | -1 | 0.28 | -0.28 | -1 | -1 | 1 | -0.28 | | |
| **Q8BJU0-2** | Sgta | Isoform 2 of Small glutamine-rich tetratricopeptide repeat-containing protein alpha | -1 | 0.1 | -0.1 | -1 | -1 | 1 | -0.1 | | |
| **Q3V4A1** | Stk11 | Serine/threonine-protein kinase STK11 | -1 | 0.28 | -0.28 | -1 | -1 | 1 | -0.28 | Y | Antroalbol H induces nuclear translocation of Stk11 in adipocytes (Wang et al., 2019)<br>Metabolic stress induces Stk11 nuclear exit via exportin 7 in astrocytes (H. J. Liang et al., 2015)<br>Imp-α cargo. Stk11 mostly in the nucleus and the binding of Strad and Mo25 modulate its nuclear exit (Dorfman & Macara, 2008) |
| **A2AG41** | Stoml2 | Stomatin-like protein 2, mitochondrial | -1 | 0.17 | -0.17 | -1 | -1 | 1 | -0.17 | | |
| **Q5D0A4** | Stx1a | Syntaxin 1A | -1 | 0.83 | -0.83 | -1 | -1 | 1 | -0.83 | | |
| **P61264** | Stx1b | Syntaxin-1B | -1 | 0.22 | -0.22 | -1 | -1 | 1 | -0.22 | | Nucleocytoplasmic shuttling proteins, Ran dependent in neurons (Pereira et al., 2008). |
| **Q9JKK1-2** | Stx6 | Isoform 2 of Syntaxin-6 | -1 | 0.13 | -0.13 | -1 | -1 | 1 | -0.13 | | |
| **A0A140LHJ4** | Stxbp1 | Syntaxin-binding protein 1 | -1 | 0.45 | -0.45 | -1 | 0.28 | -0.28 | 1.61 | Y | Contains NLS (Sharma et al., 2005)<br>Kainate acid and glutamate stimulation induce nuclear |

| | | | | | | | | | | | |
|---|---|---|---|---|---|---|---|---|---|---|---|
| | | | | | | | | | | | exit in neurons (Wan et al., 2010; Zhang et al., 2011) |
| **G3UZ48** | Syncrip | Heterogeneous nuclear ribonucleoprotein | -1 | 0.44 | -0.44 | -1 | 0.32 | -0.32 | 1.38 | Y | LMB traps Syncrip in cultured cortical neurons (Smart, 2017)<br>Nuclear Syncrip redistributes to cytoplasmic stress granules following stressors such as PMA, thapsigargin, arsenite and heat shock (Quaresma et al., 2009) |
| **W0TYI4** | Tcp1 | T-complex protein 1 | -1 | 0.18 | -0.18 | -1 | -1 | 1 | -0.18 | | |
| **Q3TIZ0** | Tuba1c | Tubulin alpha chain | -1 | 4.85 | -4.85 | -1 | -1 | 1 | -4.85 | | |
| **A0A0A6YVV3** | Txndc9 | Thioredoxin domain-containing protein 9 | -1 | 0.44 | -0.44 | -1 | -1 | 1 | -0.44 | | |
| **Q6ZPJ3** | Ube2o | (E3-independent) E2 ubiquitin-conjugating enzyme UBE2O | -1 | 0.44 | -0.44 | -1 | -1 | 1 | -0.44 | | Interacts with several Imp-β members with IPO7 achieving the highest score (Mashtalir et al., 2014) |
| **B2KF55** | Ube2v2 | Ubiquitin-conjugating enzyme E2 variant 2 | -1 | 0.28 | -0.28 | -1 | 0.24 | -0.24 | 1.17 | | |
| **Q99PL6-2** | Ubxn6 | Isoform 2 of UBX domain-containing protein 6 | -1 | 0.44 | -0.44 | -1 | -1 | 1 | -0.44 | | |
| **O08614** | Utrn | Utrophin | -1 | 0.44 | -0.44 | -1 | -1 | 1 | -0.44 | | |
| **D3YXN6** | Ywhaz | 14-3-3 protein zeta/delta | -1 | 0.28 | -0.28 | -1 | -1 | 1 | -0.28 | **Y** | Forskolin increases RACK1 binding with Ywhaz, essential for RACK1 nuclear import and PKA-dependent BDNF transcription. cAMP/PKA mediated nuclear accumulation of Ywhaz in SHSY5Y cells (Neasta et al., 2012)<br>Interacts with Imp-α2 and Imp-α4 (Arjomand et al., 2014) |
